# Antigen-presenting type-I conventional dendritic cells facilitate curative checkpoint blockade immunotherapy in pancreatic cancer

**DOI:** 10.1101/2023.03.05.531191

**Authors:** Krishnan K. Mahadevan, Allison M. Dyevoich, Yang Chen, Bingrui Li, Hikaru Sugimoto, Amari M. Sockwell, Kathleen M. McAndrews, Huamin Wang, Shabnam Shalapour, Stephanie S. Watowich, Raghu Kalluri

**Author notes:** Corresponding author: Raghu Kalluri, MD, PhD.

## Abstract

Inflammation and tissue damage associated with pancreatitis can precede or occur concurrently with pancreatic ductal adenocarcinoma (PDAC). We demonstrate that in PDAC coupled with pancreatitis (ptPDAC), antigen-presenting type-I conventional dendritic cells (cDC1s) are specifically activated. Immune checkpoint blockade therapy (iCBT) leads to cytotoxic CD8^+^ T cell activation and eradication of ptPDAC with restoration of lifespan even upon PDAC re-challenge. Such eradication of ptPDAC was reversed following specific depletion of dendritic cells. Employing PDAC antigen-loaded cDC1s as a vaccine, immunotherapy-resistant PDAC was rendered sensitive to iCBT with a curative outcome. Analysis of the T-cell receptor (TCR) sequences in the tumor infiltrating CD8^+^ T cells following cDC1 vaccination coupled with iCBT identified unique CDR3 sequences with potential therapeutic significance. Our findings identify a fundamental difference in the immune microenvironment and adaptive immune response in PDAC concurrent with, or without pancreatitis, and provides a rationale for combining cDC1 vaccination with iCBT as a potential treatment option.

## Introduction

The link between cancers and inflammation was first made by Rudolph Virchow more than a century ago when cancers were observed at sites with chronic inflammation (*1*). The host response to cancer shares many parallels with inflammation and wound healing process (*1-3*). Epidemiological data indicates that patients with acute or chronic pancreatitis (CP) have an increased risk of developing PDAC (*4-6*). Although CP is a rare disease with an incidence rate of 7-10 per 100,000 persons (*7*), a substantial portion (nearly 9%) of PDAC patients are documented with a previous history of CP (*8*). In addition to PDAC arising in patients with pre-existing chronic pancreatitis (*5*), pancreatitis frequently accompanies the progression of PDAC due to bile duct obstruction as a result of external compression from the tumor or due to tumor invasion into the biliary tree (*9*). Although pancreatitis has been shown to increase the risk of PDAC (*10, 11*) (*5*), murine models demonstrate that pancreatitis by itself failed to initiate PDAC in the absence of driver mutations (*12*). Therefore, how pancreatitis influences PDAC progression, the immune microenvironment, and adaptive immune response remains unknown.

PDAC is among the cancers that remains refractory to immunotherapy, as demonstrated by murine models and clinical trials (*13-15*). Several reasons have been attributed to the resistant nature of PDAC to immunotherapy, including a lack of effector T cell response (*16*), paucity of neo-antigens, lack of antigen-presenting cells (*17, 18*), and presence of suppressive myeloid and regulatory T cells (Tregs) in the TME (*19, 20*). Studies that probe the functional role of T cells and its subsets arrive at contradictory conclusions in autochthonous model of PDAC, pancreatitis associated PDAC (ptPDAC) models and orthotopic models of PDAC (*17, 21, 22*) (*23, 24*). However, the influence of accompanying pancreatitis, its impact on antigen presentation and T cell immunity remains to be addressed. Therefore, understanding the functional role of inflammation in modifying the T cell response in PDAC will offer valuable insights into how best exploit the potential of the immunotherapy in treatment of PDAC.

In this study, we determine the impact of pancreatitis on the immune microenvironment of PDAC and whether pancreatitis alters the functional contribution of CD4^+^ or CD8^+^ T cells in PDAC progression. Our study unravels fundamental differences in the tumors with and without accompanying pancreatitis during PDAC initiation. Our study identifies the emergence of activated dendritic cells in pancreatitis and highlights the importance of cDC1s in sensitizing PDAC to checkpoint blockade immunotherapy. Further, we explore whether PDAC antigen-loaded cDC1s could be of therapeutic benefit in combination with immunotherapy to eradicate PDAC.

## Results

### Pancreatitis associated inflammation recruits dendritic cells and myeloid cells with repression of T cells in wild type mice

To understand the impact of pancreatitis on the immune infiltrates, we performed flow cytometry and CyTOF analysis on the pancreas of WT mice after induction short-term and long-term pancreatitis (SP and LP respectively) **(Figure 1A)**. We induced pancreatitis with 4 injections per day of caerulein (an analogue of cholecystokinin that is implicated in the pathogenesis of pancreatitis) on two days, 24 hours apart in WT mice. This was defined as short-term pancreatitis. Multiple bouts of acute pancreatitis (three days per week) with caerulein injections over 2-3 weeks was defined as long-term pancreatitis (*25*) **(Figure 1A)**. Induction of SP resulted in tissue edema, and an increase in the number of lymphoid structures (LS) **(Supplementary figure S1A-S1D)**. In WT mice with SP, FlowSOM analysis of CyTOF data with unsupervised hierarchical clustering of CD45^+^ cells identified 18 MCs that were grouped into B cells (metacluster (MC) 1-2), T cells (MC 3-6), DCs (MC 7-10) and myeloid (MC 11-17) populations **(Figure 1B-1C)**. Among the T and B cell populations, SP resulted in a decrease in frequencies of CD40^+^CD19^+^ B cells (MC1) **(Figure 1C-1D)**, and in CD4^-^CD8^-^ negative T cells (CD3^+^ T - MC3; CD3^+^Ly-6C^+^ - MC6), and CD3^+^CD8^+^ T cells (MC5) **(Figure 1C and 1E)**. Among the DC populations, SP resulted in an increase in the frequencies of CD45^+^CD11c^+^ DCs (MC8) **(Figure 1C and 1F)**. For separation of CD11c^+^ cells into DC and myeloid MCs, F4/80^-^Ly-6G^-^CD11c^+^ cells were identified as DCs and F4/80^+^ or Ly-6G^+^CD11c^+^ cells were grouped under myeloid MCs. Further, SP resulted in an increase in the frequencies of CD11b^+^F4/80^+^CD11c^+^ (MC11); CD11b^+^Ly-6G^hi^CD11c^+^(MC12); CD11b^+^Ly-6G^lo^Ly-6C^hi^ (MC14); CD11b^+^Ly-6C^hi^Ly-6G^lo^PD-L1^+^ (MC15) and CD11b^+^F4/80^+^Ly-6G^lo^ (MC16) myeloid populations **(Figure 1C and 1G).** However, a decrease in proportion of one of the minor DC populations CD11b^+^CD80^+^CD11c^+^ CD40^+^PD-L1^+^ (MC10) was observed in WT-SP mice **(Figure 1F).**

**Figure 1:**
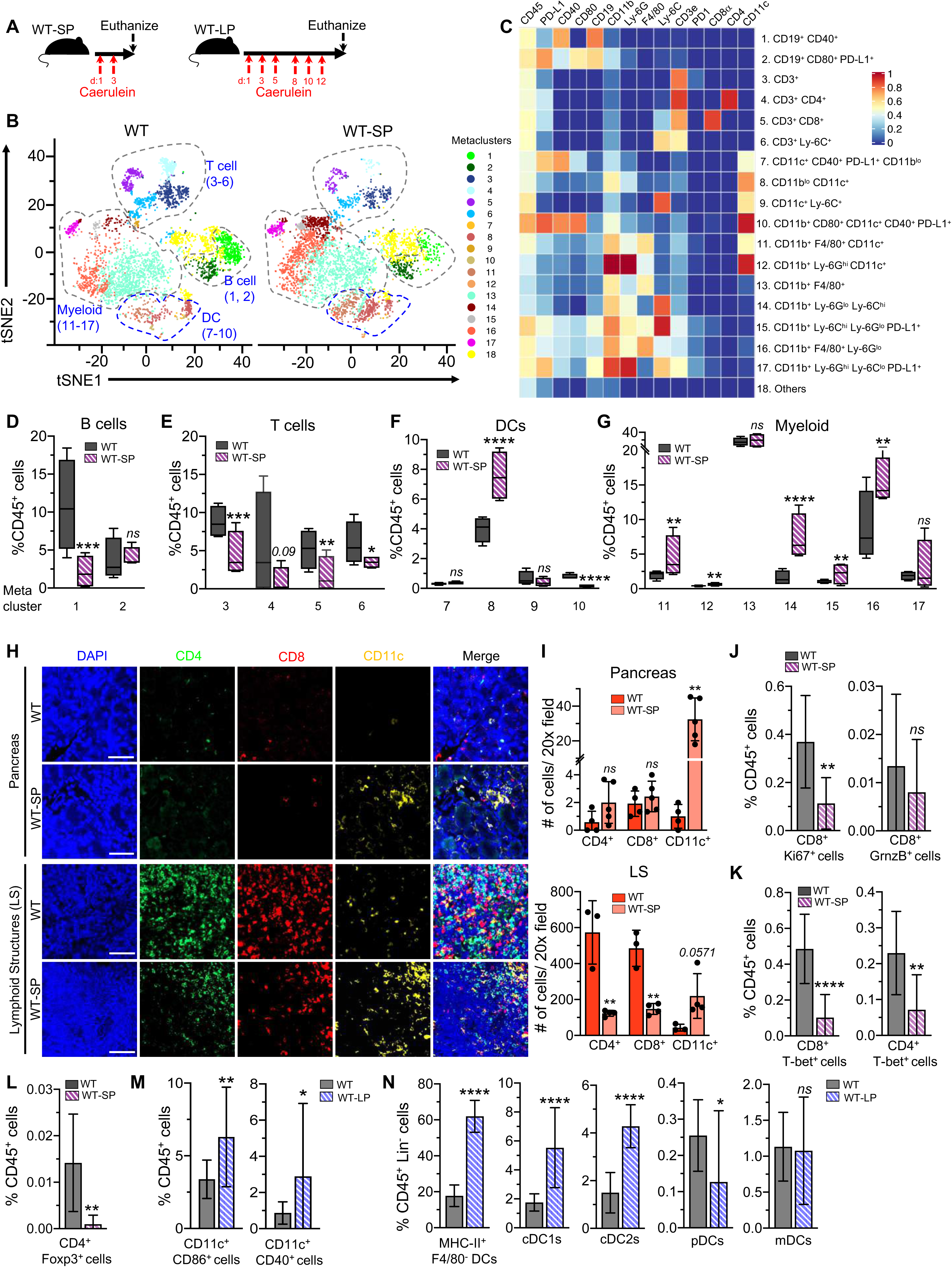
Pancreatitis recruits activated dendritic cells and suppresses T cells in the pancreas of WT mice. (**A**) Schematic representation of SP and LP induction in WT mice. (**B**) Representative viSNE plots on CD45^+^ cells of WT (n=16 mice, pancreata from 4 mice combined per sample) and WT-SP mice pancreas (n=8 mice, pancreata from 2 mice combined per sample) assessed by CyTOF. (**C**) Heat map of WT pancreas infiltrating immune cell metaclusters (MCs) displaying expression values of individual parameters normalized to the maximum mean value across MCs. (**D-G**) Relative frequencies of MCs by unsupervised clustering of CyTOF markers in indicated groups: B cell MCs (**D**), T cell MCs (**E**), DC MCs (**F**), and myeloid cell MCs (**G**). (**H-I**) Representative CD4, CD8, CD11c immunostaining (**H**), with quantification of pancreas and LS in pancreas (**I**), of WT and WT-SP mice (n=3-5/ group). (**J-L**) Immunophenotyping analysis of T cell populations in WT and WT-SP mice by flow cytometry. Relative frequencies of immune cells in indicated groups. CD8^+^Ki67^+^ and CD8^+^GrnzB^+^ (**J**); CD8^+^T-bet^+^ and CD4^+^, T-bet^+^ (**K**); and CD4^+^Foxp3^+^ cells (**L**) measured as a percentage of CD45^+^ cells. (**M-N**) Immune cell populations from pancreata of WT (n=20 mice, pancreata from 4 mice were combined per replicate), WT-LP (n=10 mice, pancreata from 2 mice were combined per replicate) mice evaluated by flow cytometry. (**M**) CD11c^+^CD86^+^ DCs and CD11c^+^CD40^+^ DCs were measured as a percentage of CD45^+^ cells. (**N**) CD11c^+^MHC-II^+^F4/80^+^ DCs; cDC1s (CD11c^+^B220^-^CD172a^-^CD64^-^Ly-6C^-^CD11bMHC-II^+^XCR1^+^ cells); cDC2s (CD11c^+^ B220^-^CD172a^+^CD64^-^Ly-6C^-^CD11b^+^ cells); pDCs (CD11c^+^B220^+^SIGLEC H^+^ cells) and mDCs (CD11c^+^B220^-^CD172a^+^CD64^-^ Ly-6C^+^ cells) were measured as a percentage of CD45^+^Lin^-^ (CD19^-^Ly-6G^-^CD3^-^NK1.1^-^) cells. In **D-G,** data are presented as box and whisker plots (min to max) and as mean ± SD in **I, J, K, L, M,** and **N**. Significance was determined by unpaired t-test in **D, E, F, G, J, K, L, M** and **N**. In **I**, Mann-Whitney test was used for comparison of CD11c^+^ cells in the LS and unpaired T-test was used for all other comparisons. *P <0.05, **P <0.01, ***P <0.001, ****P <0.0001, *ns*: not significant. Scale bars indicate 100 μm. See related supplementary figures S1-S2.

Increased CD11c^+^ and CD11b^+^ cells in pancreatic acini and LS in SP was confirmed by immunostaining **(Figure 1H-1I, Supplementary figure S1E-S1F)**. A decrease in both CD4^+^ and CD8^+^ T cells were seen in LS, in contrast to pancreatic acini where insignificant differences in CD4^+^ or CD8^+^ T cells were observed in WT-SP mice (**Figure 1H-1I).** Further analysis of T cell populations by flow cytometry revealed global decrease in frequencies of proliferating Ki-67^+^ and T-bet^+^CD8^+^ T cells; Th_1_ cells and Tregs in WT-SP mice **(Figure 1J-1L, Supplementary figure S1G)**. Next, we analyzed the immune infiltration from the WT vs. WT-LP mice. We confirmed induction of pancreatitis via measurement of serum amylase and lipase and observed increased concentrations of both enzymes in WT-LP mice compared to controls **(Supplementary Figure S2A)**. Analysis of dendritic cells in WT-LP pancreas demonstrated increase in frequency of activated CD86^+^ and CD40^+^CD11c^+^DCs, MHC-II^+^F4/80^-^DCs, cDC1s and other DC subsets **(Figure 1M and 1N)**. WT-LP mice phenocopied the histological and immunological findings observed in the WT-SP mice **(Supplementary Figure S2B-S2N, S1A-S1G, and Figure 1H-1N)**.

### Pancreatitis accelerates PDAC associated with recruitment of activated dendritic cells and type-I conventional dendritic cells

To analyze the impact of pancreatitis on tumor initiation, we induced SP in 7 weeks old KC (*Pdx1-Cre; LSL-Kras^G12D/+^*) mice. The KC-SP mice were euthanized 4 days after the final injection to perform CyTOF analysis of the tumor infiltrating CD45^+^ cells **(Supplementary Figure S3A)**, and again after 3 weeks to conduct histological immunostaining analysis **(Figure 2A)**. Age matched KC mice without SP (KC-10w) and disease stage-matched KC mice (KC-25w) were used as controls. In KC mice with SP, FlowSOM analysis of CyTOF data with unsupervised hierarchical clustering of CD45^+^ cells identified 12 metaclusters encompassing B cells (MC 1), T cells (MC 2,3), DCs (MC 5), and myeloid cells (MC 4, 6-11) **(Supplementary Figure S3B-S3G)**. SP in KC mice resulted in an increase in the proportion of CD45^+^, CD11c^+^ cells (MC5), while insignificant differences in B cell and T cell MCs were noted **(Supplementary Figure S3D-S3F).** SP also resulted in an increase in CD11b^+^F4/80^+^ cells (MC 10) **(Supplementary Figure S3C and S3G**). Immunophenotyping analysis by Boolean gating of the CyTOF data also confirmed that SP in KC mice resulted in an increase in DCs within the immune cell population (CD45^+^CD11c^+^ cells) **(Supplementary Figure S3H)**.

**Figure 2:**
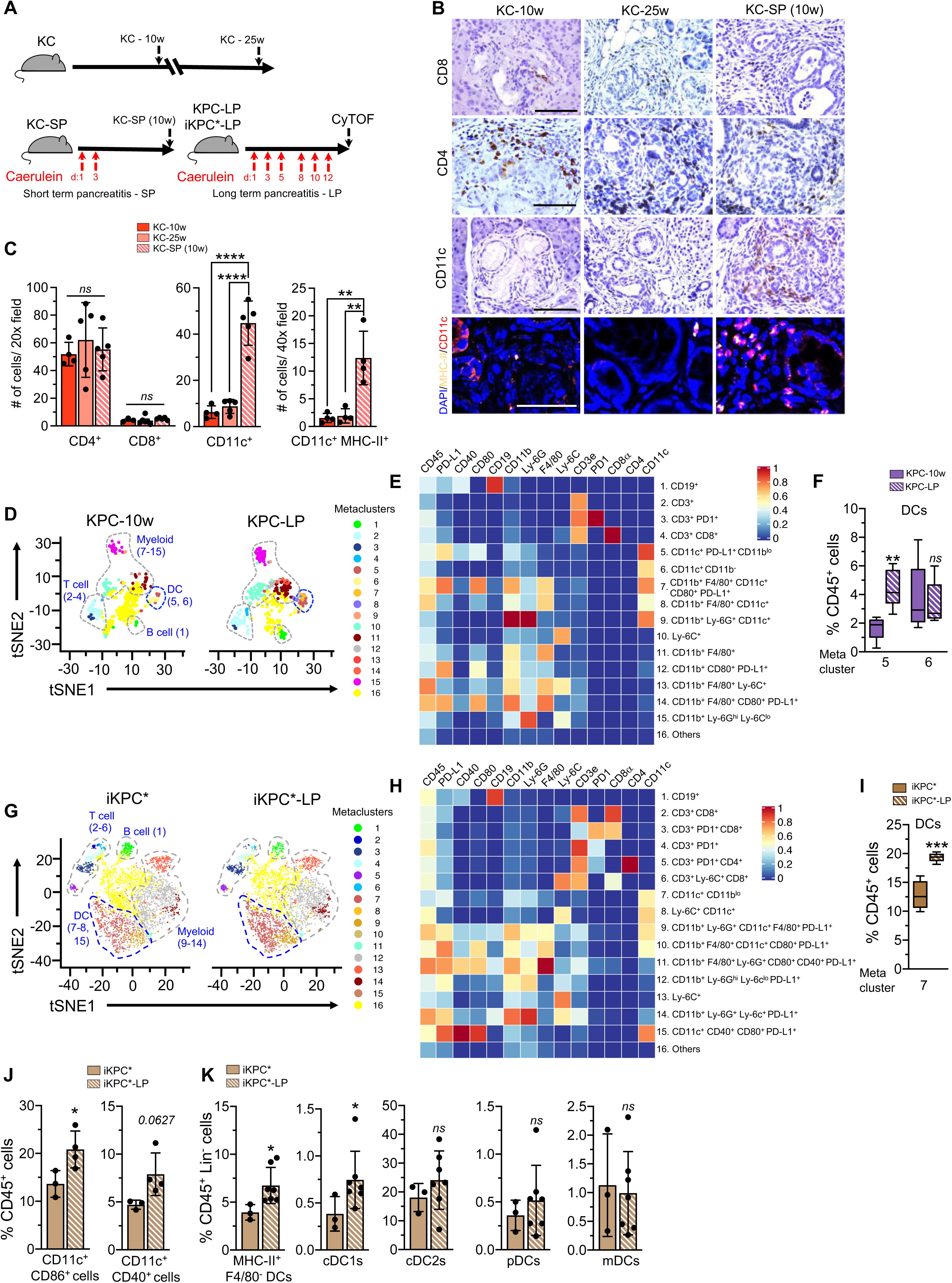
Pancreatitis recruits activated dendritic cells and cDC1s to the PDAC TME. (**A**) Schematic representation of induction of short-term pancreatitis (SP) (each red arrow indicates injection of caerulein 4 times per day, 6 hours apart) in KC mice with age and stage matched controls and induction of long-term pancreatitis (LP) in age matched KPC and orthotopic iKPC* mice. (**B-C**) Representative CD4, CD8, CD11c and CD11c-MHC-II immunostaining (**B**), with quantification of PanIN lesions (**C**). KC-10w (n=3-4), KC-25w (n=4-6) and KC-SP (10w) (n=4-5). (**D**) Representative viSNE plots of CD45^+^ cells in KPC-10w (n=5) and KPC-LP (n=6) mice pancreata assessed by CyTOF. (**E**) Heat map of KPC mice pancreata infiltrating immune cell metaclusters (MCs) displaying expression values of individual parameters normalized to the maximum mean value across MCs. (**F**) Relative frequencies of DC MCs by unsupervised clustering of CyTOF markers. (**G**) Representative viSNE plots on CD45^+^ cells of iKPC* (n=4) and iKPC*-LP (n=6) mice pancreata assessed by CyTOF. (**H**) Heat map of iKPC* pancreata infiltrating immune cell MCs displaying expression values of individual parameters normalized to the maximum mean value across MCs. (**I**) Relative frequencies of DC MCs of indicated experimental groups measured as a percentage of CD45^+^ cells. (**J-K**) Immune cell populations from pancreata of iKPC* (n=3), iKPC*-LP (n=4-7) mice evaluated by flow cytometry. (**J**) CD11c^+^, CD86^+^ DCs and CD11c^+^, CD40^+^ DCs were measured as a percentage of CD45^+^ cells. (**K**) CD11c^+^MHC-II^+^F4/80^+^ DCs; cDC1s (CD11c^+^B220^-^CD172a^-^CD64^-^Ly-6C^-^, CD11b^-^MHC-II^+^XCR1^+^ cells); cDC2s (CD11c^+^B220^-^CD172a^+^CD64^-^Ly-6C^-^CD11b^+^ cells); pDCs (CD11c^+^B220^+^SIGLEC H^+^ cells) and mDCs (CD11c^+^B220^-^CD172a^+^CD64^-^Ly-6C^+^ cells) were measured as a percentage of CD45^+^Lin^-^ (CD19, Ly-6G, CD3, NK1.1) cells. In **F and I,** data are presented as box and whisker plots (min to max) and as mean ± SD in **C, J and K**. Significance was determined by one-way ANOVA using Dunnett’s multiple comparisons test in **C**, and by unpaired t-test in **F, I and J.** In **K**, Mann-Whitney test was used for comparison of MHC-II^+^F4/80^-^ DCs, cDC1s and unpaired T-test was used for comparison of cDC2s, mDCs and pDCs. * P<0.05, ** P<0.01, *** P<0.001, **** P<0.0001, *ns*: not significant. Scale bars indicate 100 μm. See related supplementary figures S3-S7.

Consistent with our findings from CyTOF analysis, immunostaining confirmed that SP did not impact CD4^+^ and CD8^+^ T cells infiltration in PanIN lesions and within the associated lymphoid structures (LS) in the pancreas of KC mice (**Figure 2B-2C and Supplementary Figure S4A-S4B**). Further analysis of CD4 subsets did not reveal differences in the CD4^+^Foxp3^+^ Tregs; CD4^+^GATA3^+^ T cells (Th_2_); CD4^+^T-bet^+^ T cells (Th_1_) and CD4^+^RoRγt^+^ T cells (Th_17_) between disease stage-matched KC and KC-SP mice (**Supplementary Figure S4C-S4D**). KC-SP mice showed an increase in the number of CD11c^+^ DCs in the PanIN lesions quantified by immunostaining **(Figure 2B-2C).** The LS harbored the majority of the baseline CD11c^+^ cells in the KC mice **(Supplementary Figure S5A-S5B)**, with PanIN lesions revealing minimal CD11c^+^ cells. SP in KC mice resulted in a significant increase in CD11c^+^ DCs in PanIN lesions, whereas only a modest increase was observed in the LS **(Figure 2C and Supplementary Figure S5A-S5B)**. Analysis of the myeloid marker, CD11b, revealed insignificant differences in the PanIN lesions of KC-SP mice (**Supplementary Figure S5C and S5D**). Analysis of the activation marker, MHC-II on CD11c^+^ DCs, revealed that SP significantly recruited MHC-II^+^, CD11c^+^ DCs into the PanINs of KC mice **(Figure 2B-2C and Supplementary Figure S5E)**. SP resulted in acceleration of tumor initiation in the KC mice as measured by histological presence of ADM/PanIN lesions **(Supplementary Figure S6A and S6B)**. Moreover, we observed an increase in the number of LS in pancreas as a response to long-term pancreatitis (LP (**Supplementary Figure S6C-S6E**).

Next, we analyzed the immune infiltration in KPC (*Pdx1-Cre, LSL-Kras^G12D/+^*, *Trp53^R172H/+^*) and the highly aggressive orthotopic iKPC* mice (orthotopically implanted PDAC cell lines with *P48-Cre*; *tetO-LSL-Kras^G12D/+^; Trp53^L/L^* genotype) with LP (**Figure 2A**). LP accelerated tumor growth in the KPC mice **(Supplementary Figure S7A)**. Similar to KC mice with pancreatitis, LP in KPC (KPC-LP) and iKPC* (iKPC-LP) mice resulted in an increase in frequencies of DC subpopulations as following: CD11c^+^PDL1^+^ DCs (MC 5) in KPC mice and CD11c^+^CD11b^lo^ in iKPC* mice (MC 7) (**Figure 2D-2I**) with insignificant changes in the frequencies of B and T cells among the immune cell population (MC1-4) **(Supplementary Figure S7B-S7I)**. Among the myeloid MCs, a significant increase in the percentages of CD11c^+^CD11b^+^F4/80^+^PDL1^+^CD80^+^ cells (MC7); CD11b^+^F4/80^+^CD11c^+^ cells (MC8); CD11b^+^CD80^+^PDL1^+^ cells (MC12); CD11b^+^F4/80^+^CD80^+^PDL1^+^ cells (MC14) and CD11b^+^Ly-6G^hi^Ly-6C^lo^ cells (MC15) was observed in KPC mice **(Figure 2E, Supplementary Figure S7C and S7D),** whereas insignificant differences in the myeloid MCs were observed in the orthotopic iKPC* mice **(Supplementary Figure S7H and S7I)**. Further analysis of activation markers such as CD86 and CD40 revealed that LP increased the proportion of CD86^+^CD11c^+^ DCs in the orthotopic iKPC* mice **(Figure 2J)**. While pancreatitis induced an increase in the percentage CD11c^+^ cells, murine macrophages can also express dendritic cell markers such as CD11c and MHC-II (*26*). We confirmed that pancreatitis resulted in an increase in frequencies of MHC-II^+^F4/80^-^CD11c^+^ cells **(Figure 2K)**. DCs are subdivided into distinct functional subsets, conventional DCs (cDCs), plasmacytoid DCs (pDCs) and monocytic DCs (mDCs) (*26-28*). Among conventional DCs, the cDC1s are important for the priming of tumor antigen specific T cell and facilitating their responses to restrain tumors, whereas cDC2s were found to be involved in tolerance induction that facilitate tumor growth (*27*). Analysis of cDC1s in iKPC* mice showed that LP resulted in an increase in frequencies of cDC1s in the PDAC TME **(Figure 2K)**.

Taken together, our results indicate that changes in adaptive immune cells, particularly T and B cells, in WT mice with pancreatitis represents a putative tolerogenic mechanism to prevent further pancreatic damage in association with self-antigen presentation by DCs during the course of inflammation (*25, 29*). Whereas in the context of tumor bearing KC, KPC and orthotopic iKPC* mice with pancreatitis, the T cell recruitment might reflect presentation of potential tumor-associated antigens by activated DCs and cDC1s.

### CD4^+^ T cells promote PDAC initiation and progression in mice with pancreatitis by restraining the function of CD11c^+^ DCs

Next, we examined the impact of CD11c^+^ DCs recruitment induced by pancreatitis on T cell function, and probed whether the cross talk between DCs and T cells is responsible for the observed acceleration of PDAC development in background of inflammation. We crossed *CD4^-/-^* or *CD8^-/-^* mice with *Pdx1-Cre, LSL-Kras^G12D/+^*, *Trp53^R172H/+^* (KPC) and generated *CD4^-/-^, Pdx1-Cre, LSL-Kras^G12D/+^* (KC CD4^-/-^); *CD4^-/-^, Pdx1-Cre, LSL-Kras^G12D/+^*, *Trp53^R172H/+^* (KPC CD4^-/-^); *CD8^-/-^, Pdx1-Cre, LSL-Kras^G12D/^* (KC CD8^-/-^) and *CD8^-/-^, Pdx1-Cre, LSL-Kras^G12D/+^*, *Trp53^R172H/+^* (KPC CD8^-/-^) mice. Lack of CD4^+^ and CD8^+^ T cells in the thymus, spleen, and tumors of KPC CD4^-/-^ and KPC CD8^-/-^ mice was confirmed by immunostaining (**Supplementary Figure S8A-S8D**). First, we analyzed tumor initiation and progression in KC mice with genetic deletion of CD4^+^ or CD8^+^ T cell populations. Age-matched analysis (at 25 weeks) of tumor initiation in the KC, KC CD4^-/-^ and KC CD8^-/-^ mice was conducted. Analysis of PanIN lesions in these mice showed insignificant differences in tumor initiation or progression (**Figure 3A-3C**). Next, to evaluate the influence of pancreatitis on pancreatic cancer initiation and progression, SP and LP were induced in 7-week-old KC, KC CD4^-/-^ and KC CD8^-/-^ mice and euthanized after 3 weeks (SP) and 8 weeks (LP) to assess cancer initiation and progression (**Figure 3A, Supplementary Figure S9A**). In contrast to tumor initiation in mice without pancreatitis, we observed that KC CD4^-/-^ mice with SP and LP had relatively fewer PanINs compared to KC and KC CD8^-/-^ mice (**Figure 3B-3E, Supplementary Figure S9A-S9C**). Since the majority of the CD4 infiltrates in the PanIN lesions were Tregs, Th_2_ and Th_17_, which have been previously found to support tumor development **(Supplementary Figure S4C and S4D)**, we hypothesized that the CD4^+^ T cells regulate the tumor-directed cytotoxic CD8^+^ T lymphocyte (CTL) response which results in inhibition of tumor progression in KC-SP mice. Depletion of CD8^+^ T cells with αCD8 antibody treatment resulted in re-emergence of PanIN lesions in the KC-SP CD4^-/-^ mice **(Figure 3D and 3E, Supplementary Figure S9D and S9E)** suggesting that CD4^+^ T cells promote tumorigenesis specifically in KC mice with pancreatitis via inhibition of the CTL response.

**Figure 3:**
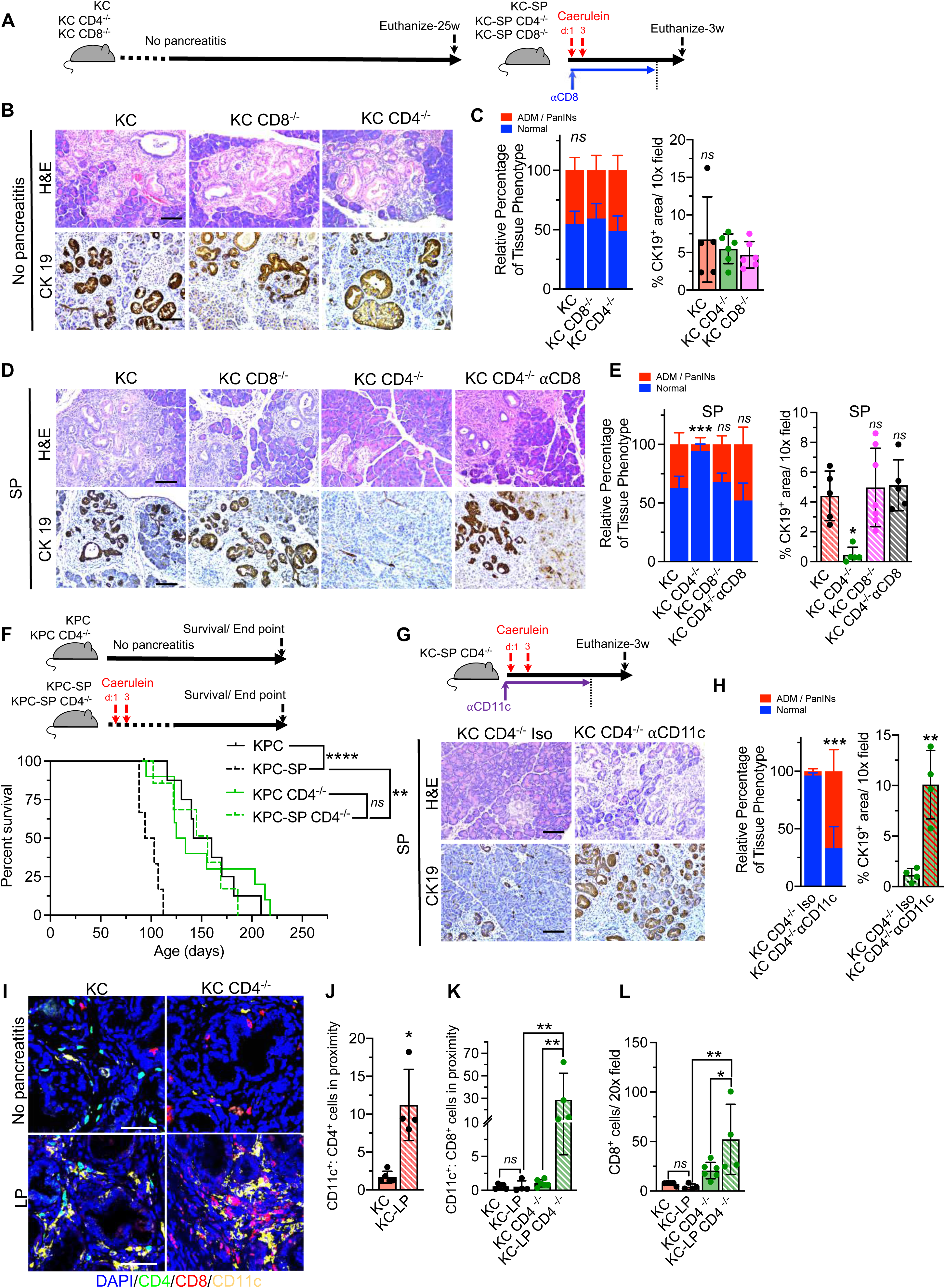
CD4^+^ T cells promote PDAC progression in KC/KPC mice with pancreatitis by restraining dendritic cells. (**A**) Schematic representation of SP induction (each red arrow indicates injection of caerulein 4 times per day, 6 hours apart) in KC, KC CD4^-/-^ and KC CD8^-/-^ mice starting from 7w of age. (**B-C**) Representative H&E and CK19 immunostaining images (**B**), with quantification of PanIN lesions based on H&E (**C**, left panel) and CK19 staining (**C**, right panel) of KC (n=5), KC CD4^-/-^ (n=6-7) and KC CD8^-/-^ (n=6) mice pancreata. (**D-E**) Representative H&E and CK19 immunostaining (**D**), with quantification of PanIN lesions (**E**), of KC-SP (n=5), KC-SP CD4^-/-^ (n=5), KC-SP CD8^-/-^ (n=6-7), KC-SP CD4^-/-^ CD8 (n=5) mice pancreata. Note: Same KC-SP control mice from Figure 2B. (**F**) Schematic representation of SP induction in KPC and KPC CD4^-/-^ mice starting at 6w of age (top panel) and Kaplan-Meier survival curves of KPC (n=9), KPC-SP (n=6), KPC CD4^-/-^ (n=10), KPC-SP CD4^-/-^ (n=7) (bottom panel). (**G**) Schematic representation of SP induction in KC⍰ CD4^-/-^ mice with ⍰ CD11c or isotype antibody treatment time points denoted with purple arrows (top panel) and representative H&E and CK19 immunostaining of PanIN lesions (bottom panel). (**H**) Quantification of H&E and CK19 immunostaining of PanIN lesions of ⍰ CD11c or isotype treated KC-SP CD4^-/-^ mice (n=4/ group). (**I-L**) Representative images of CD4, CD8 and CD11c co-staining (**I**), with quantification of CD11c^+^ and CD4^+^ cells in spatial proximity (**J**), CD11c^+^ and CD8^+^ cells in spatial proximity (**K**), and CD8^+^ cells (**L**), in KC and KC CD4^-/-^ mice with and without LP. KC (n=5), KC-LP (n=4), KC CD4^-/-^ (n=6) and KC-LP CD4^-/-^ (n=4). In **C, E, H, J, K and L** data are presented as mean ± SD. Significance was determined by two-way ANOVA with Tukey’s test for comparison of relative percentage of tissue phenotype in **C, E** and **H**, one-way ANOVA with Dunnett’s multiple comparisons test for immunostaining in **C**, Kruskal-Wallis with Dunn’s multiple comparisons test for immunostaining in **E**, log rank test in **F**, unpaired T-test in **J** and for immunostaining in **H**, and one-way ANOVA with Sidak’s multiple comparisons test in **K** and **L**. * P <0.05, ** P <0.01, *** P <0.001, *ns*: not significant. Scale bars indicate 100 μm. See related supplementary figures S8-S10.

Next, to determine whether CD4^+^ T cells accelerate PDAC in the KPC mice with pancreatitis as well, we induced SP in 6w old KPC and KPC CD4^-/-^ mice **(Figure 3F)**. The KPC and KPC CD4^-/-^ mice demonstrated similar overall survival kinetics **(Figure 3F)**. SP accelerated PDAC progression in the KPC mice (KPC-SP) **(Figure 3F)**. The KPC-SP mice succumbed earlier to PDAC and demonstrated shorter overall survival when compared to KPC-SP CD4^-/-^ mice **(Figure 3F)** confirming our results with the KC mice. Histological analysis at end point showed similar levels of invasive PDAC in all groups confirming that these mice reached endpoints due to PDAC and not from potential pancreatitis related morbidity **(Supplementary Figure S9F and S9G)**. Therefore, while CD4^+^ T cells had no impact on the progression of PanINs and PDAC in KC and KPC mice respectively, they promoted PanINs and PDAC in KC and KPC mice overlaid with pancreatitis.

Further, to understand the functional crosstalk between CD4^+^ T cells and CD11c^+^ DCs in KC mice with pancreatitis, we treated KC-SP CD4^-/-^ mice with αCD11c antibody to deplete CD11c^+^ DCs (**Figure 3G, Supplementary Figure S9H and S9I**). Depletion of CD11c^+^ DCs in KC-SP CD4^-/-^ mice restored PanIN formation (**Figure 3G-3H**), indicating that CD4^+^ T cells promote tumor initiation either by inhibition of CD11c^+^DCs or CD8^+^ T cells to promote tumor initiation. Although KC-LP demonstrated an increase in CD11c^+^ DCs **(Figure 3I and Supplementary Figure S10A-S10D)** without alterations in the number of T cells associated with PanIN lesions, KC-LP mice showed increased spatial proximity of CD11c^+^ DCs to CD4^+^ T cells **(Figure 3I-3J and Supplementary Figure S10A)**, which predominantly consisted of the inhibitory T cell subsets. Lack of CD4^+^ T cells removed their inhibitory effect on anti-tumor directed response, increasing CD11c^+^ DC: CD8^+^ T cell proximity, likely contributing to PanIN inhibition in the presence of pancreatitis (**Figure 3I and 3K-3L**). Overall, our results indicate that pancreatitis recruited activated DCs into pancreas of KC mice. Despite an increase in the CD11c^+^ DCs, inhibitory CD4^+^ T cells override the anti-tumorigenic effect of DCs in KC-SP/LP setting. A critical step in the development of T cell response involves contact between the CD8^+^ T cell and the antigen presenting cells (*30*). Lack of CD4^+^ T cells in KC-SP CD4^-/-^ mice likely enabled the functionality of activated DCs via engagement of CD8^+^ T cells, resulting in inhibition of PanIN formation and progression.

### Pancreatitis sensitizes PDAC to curative immune checkpoint blockade therapy

We hypothesized that the increased infiltration of activated DCs and their spatial proximity to T cells in the presence of pancreatitis may render pancreatic cancer sensitive to checkpoint blockade in this setting. Multiple studies in PDAC patients and in pre-clinical models have established the failure of immune checkpoint blockade therapy (iCBT) in PDAC (*13-15*). We investigate whether combination of antigen presenting cDC1s with αPD1 or αCTLA4+αPD1 iCBT might inhibit PanIN initiation and PDAC progression (**Figure 4A**). KC-SP mice treated with αPD1 or αCTLA4+αPD1 demonstrated inhibition of tumor initiation compared to the isotype control mice and untreated KC mice (**Figure 4B and 4C**). Our results showed that pancreatitis associated KC (KC-SP) mice were sensitive to checkpoint blockade, further validating our findings with KC-SP CD4^-/-^ mice (vide supra, **Figure 3D and 3G**).

**Figure 4:**
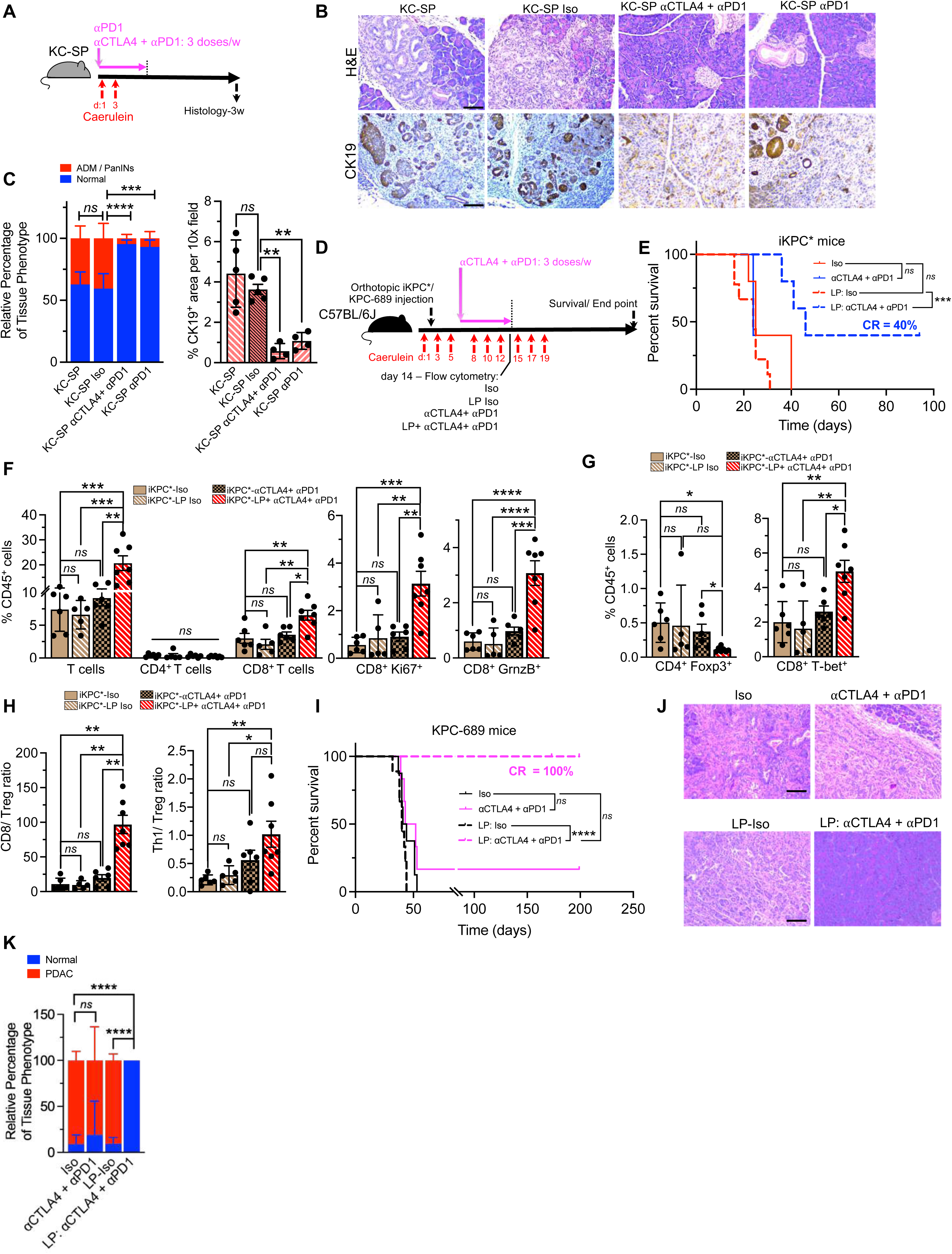
Pancreatitis sensitizes PanIN lesions in KC mice and orthotopic iKPC* tumors to checkpoint blockade immunotherapy. (**A**) Schematic representation of SP induction (each red arrow indicates injection of caerulein 4 times per day, 6 hours apart) and isotype antibody and ⍰PD1, ⍰CTLA4 + ⍰PD1 treatment in KC mice. (**B-C)** Representative H&E and CK19 immunostaining (**B**), with quantification of PanIN lesions by H&E (**C**, left panel) and CK19 staining (**C**, right panel) of isotype, ⍰CTLA4 + ⍰PD1, ⍰PD1 treated mice. KC-SP (n=5), KC-SP Isotype (n=5), KC-SP ⍰PD1 (n=4) and KC-SP ⍰CTLA4 + ⍰PD1 (n=4). Note: Same KC-SP control mice from Figure 2B. (**D**) Schematic representation of orthotopic injection of KPC-689 GFP-Luc cells or iKPC* cell line in C57BL/6J mice, LP induction (each red arrow indicates injection of caerulein 4 times per day, 6 hours apart) with ⍰CTLA4 + ⍰PD1 or isotype treatment. Analysis of tumor infiltrating lymphocytes of iKPC* mice was performed by flow cytometry in Isotype (n=6), LP: Isotype (n=5), ⍰CTLA4 + ⍰PD1 (n=6), and LP + ⍰CTLA4 + ⍰PD1 (n=7) treated iKPC* mice on day 14 (1 week following the administration of last dose of DC vaccine). (**E**) Kaplan-Meier survival curve of iKPC* mice treated with isotype (n=5), ⍰CTLA4 + ⍰PD1 (n=5), LP: isotype (n=9), and LP: ⍰CTLA4 + ⍰PD1 (n=5). (**F-L**) Immunophenotyping data by flow cytometry of indicated groups. CD3^+^(T cells); CD3^+^CD4^+^ (CD4^+^ T cells); CD3^+^CD8^+^ (CD8^+^ T cells); CD8^+^Ki67^+^ and CD8^+^GrnzB^+^ cells (**F**); CD4^+^Foxp3^+^ and CD8^+^T-bet^+^ cells (**G**) were measured as a percentage of CD45^+^ cells. CD8/Treg ratio and Th_1_/Treg ratio (**H**). (**I**) Kaplan-Meier survival curve of indicated groups. KPC-689 tumors with isotype (n=8), ⍰CTLA4 + ⍰PD1 (n=6), LP: Iso (n=9), LP: ⍰CTLA4 + ⍰PD1 (n=8), and KPC-689 parental tumors (n=8). (**J-K**) Representative H&E (**J**), and histological analysis (**K**), of orthotopic KPC-689 tumors or pancreata of indicated groups at autopsy (n=4-8/ group). In **C, F, G, H** and **K,** data are presented as mean ± SD. Significance was determined by two-way ANOVA with Tukey’s test for comparison of relative percentage of tissue phenotype in **C**, one-way ANOVA with Dunnett’s multiple comparisons test for immunostaining in **C**, log-rank test in **E**, Kruskal-Wallis with Dunn’s multiple comparisons test for comparison of CD4^+^ T cells in **F** and one-way ANOVA with Sidak’s multiple comparisons test for all other comparisons in **F**, Kruskal-Wallis with Dunn’s multiple comparisons test for comparison of CD4^+^ Foxp3^+^ T cells in **G** and one-way ANOVA with Sidak’s multiple comparisons test for all other comparisons in **G**, Kruskal-Wallis with Dunn’s multiple comparisons test for comparison of CD8/ T reg ratio in **H** and one-way ANOVA with Sidak’s multiple comparisons test for all other comparisons in **H**. * P <0.05, ** P <0.01, *** P <0.001, **** P <0.0001, *ns*: not significant. Scale bars indicate 100 μm. See related supplementary figures S11-S12. Note: KC-SP mice used in Fig. 4B and 4C are repeated from Fig. 3D and 3E respectively to allow for direct comparisons.

To examine whether inflammation recruited cDC1s can sensitize PDAC to αCTLA4+αPD1 iCBT, we induced LP in orthotopic iKPC* and KPC-689 tumor bearing mice. Six to eight weeks old C57BL/6J mice with or without LP were orthotopically injected with iKPC* or KPC-689 GFP-Luc cells (*Pdx1-Cre; LSL-Kras^G12D/+^; P53^R172H/+^* cells expressing GFP-Luc) and treated with αCTLA4+αPD1 iCBT **(Figure 4D).** The iKPC* mice did not respond to αCTLA4+αPD1 iCBT, whereas LP associated iKPC* mice demonstrated longer survival with cures in response to αCTLA4+αPD1 iCBT (**Figure 4E, Supplementary Figure S11A and S11B**). LP by itself did not alter the progression in orthotopic tumor models (**Figure 4E-4K and Supplementary Figure S11A-S11E**). Analysis of the T cell response to αCTLA4+αPD1 iCBT in LP mice revealed increased frequencies of CTLs, CD8^+^ T cell proliferation, activation and memory, decreased Tregs frequency and increase in T-bet^+^ effector CD8^+^ T cells with an increase in CD8/Treg ratio and Th_1_/ Treg ratio (**Figure 4F-H, Supplementary Figure S11F-S11I**). Consistent with the iKPC* mice, LP combined with αCTLA4+αPD1 iCBT also resulted in prolonged survival and cures in the KPC-689 orthotopic tumor bearing mice (**Figure 4I-4K, Supplementary Figure S12A-S12D**).

### PDAC antigens stimulated CD103^+^ cDC1 vaccine sensitizes PDAC to curative immune checkpoint blockade therapy

To directly examine whether cDC1s can sensitize PDAC to αCTLA4+αPD1 iCBT, we administered purified and PDAC antigen stimulated exogenous CD103^+^ cDC1s as a vaccine **(Figure 5A, Supplementary Figure S13A-S13B)**. For preparation of the cDC1 vaccine, we treated cDC1s with polyinosinic: polycytidylic acid (poly I:C) to induce maturation and concurrently exposed cDC1s to PDAC tumor lysates as a source for tumor antigen (AgS). We loaded cDC1s with either WT-LP pancreata-derived lysate as control, or iKPC*-LP tumor lysate or iKPC* cells-derived lysate (**Figure 5B**). iKPC* mice that received cDC1 vaccine stimulated with WT-LP lysate as the AgS did not synergize with αCTLA4+αPD1 iCBT, but cDC1 vaccines stimulated with iKPC*-LP PDAC lysate and iKPC*-cell lysate significantly increased survival and tumor eradication in most mice, compared to the control groups (**Figure 5B, Supplementary Figure S14A-S14E**). A substantial portion of the αCTLA4+αPD1 iCBT treated mice with cDC1 vaccine, cleared the iKPC* tumors even with two doses of the cDC1 vaccine.

**Figure 5:**
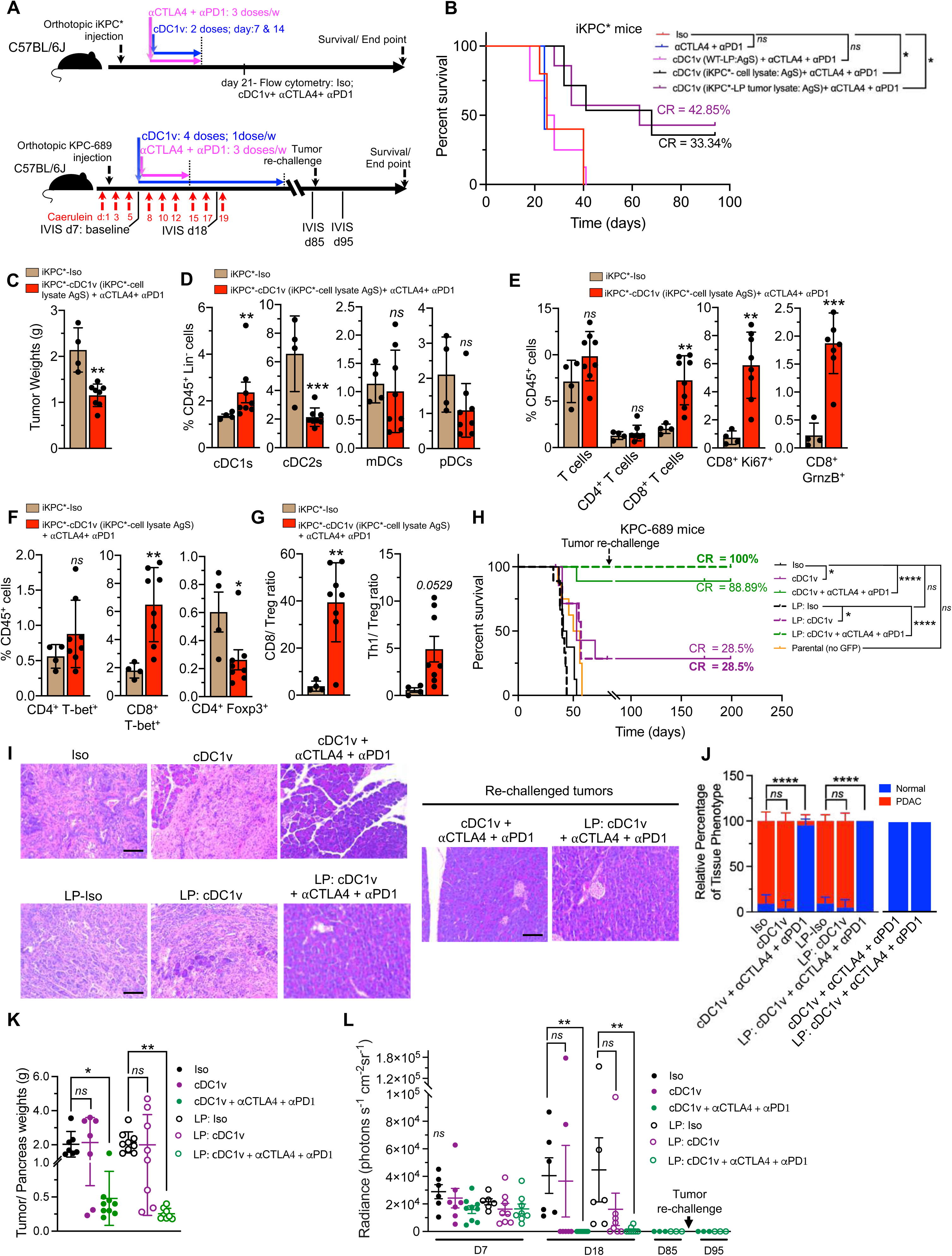
cDC1 vaccine sensitizes PDAC to curative checkpoint blockade immunotherapy. (**A**) Schematic representation of orthotopic injection of iKPC* or KPC-689 GFP-Luc cells in C57BL/6J mice, LP-induction (each red arrow indicates injection of caerulein 4 times per day, 6 hours apart), IVIS imaging and cDC1 vaccine with ⍰CTLA4 + ⍰PD1 or isotype treatment. Mice from surviving treatment groups (KPC-689 orthotopic) were re-challenged with KPC-689 cells in a second orthotopic injection (n=3/ group). (**B**) Kaplan-Meier survival curve of iKPC* mice treated with isotype (n=5), ⍰CTLA4 + ⍰PD1 (n=5), cDC1v + ⍰CTLA4 + ⍰PD1 (WT-LP lysate AgS - sorted cDC1s were loaded with lysates prepared from pancreata of WT mice with LP) (n=8), cDC1v + ⍰CTLA4 + ⍰PD1 (iKPC* cell lysate AgS) (n=6), and cDC1v + CTLA4 + PD1 (iKPC*-LP tumor lysate AgS - sorted cDC1s were loaded with lysates of orthotopic iKPC* tumor bearing mice ‘on Doxycycline’ with LP) (n=7). (**C**) Tumor weights of isotype (n=4) and cDC1v (iKPC* cell lysate AgS) + ⍰CTLA4 + ⍰PD1 (n=8) treated mice euthanized on day 21 following orthotopic iKPC* injection. (**D-G**) Immunophenotyping analysis of iKPC*-isotype (n=4) and cDC1v (iKPC* cell lysate AgS) + ⍰CTLA4 + ⍰PD1 (n=8) tumors. (**D**) Immunophenotyping analysis for DC subsets. cDC1s (CD11c^+^B220^-^CD172a^-^CD64^-^Ly-6C^-^CD11b^-^MHC-II^+^XCR1^+^cells), cDC2s (CD11c^+^B220^-^CD172a^+^CD64^-^Ly-6C^-^CD11b^+^ cells), mDCs (CD11c^+^B220^-^CD172a^+^CD64^-^Ly-6C^+^ cells) and pDCs (CD11c^+^B220^+^SIGLEC H^+^ cells) measured as a percentage of CD45^+^Lin (CD19, Ly-6G, CD3, NK1.1)^-^ cells. (**E-G**) Immunophenotyping data by flow cytometry of indicated groups. CD3^+^(T cells); CD3^+^CD4^+^ (CD4^+^ T cells); CD3^+^CD8^+^ (CD8^+^ T cells); CD8^+^Ki67^+^ and CD8^+^GrnzB^+^ cells (**E**); CD4^+^, T-bet^+^ (Th_1_) and CD8^+^, T-bet^+^ cells and CD4^+^, Foxp3^+^ cells (**F**); were measured as a percentage of CD45^+^ cells. CD8/Treg ratio and Th_1_/Treg ratio (**G**). (**H**) Kaplan-Meier survival curve of indicated groups. KPC-689 tumors with isotype (n=8), cDC1v (n=7), cDC1v+ ⍰CTLA4 + ⍰PD1 (n=9), LP: Iso (n=9), LP: cDC1v (n=8), LP: cDC1v+ ⍰CTLA4 + ⍰PD1 (n=8), KPC-689 parental tumors (n=8). (**I-J**) Representative H&E (**I**), and histological analysis (**J**), of orthotopic KPC-689 tumors or pancreata of indicated groups at autopsy (n=4-8/ group). (**K**) Tumor/ pancreas weights from indicated groups at sacrifice or endpoint. (**L**) Quantification of IVIS imaging of baseline on d7, follow up on d18, imaging on d85 before re-challenge and on d95 in indicated experimental groups. KPC-689 tumors with isotype (n=6), ⍰CTLA4 + ⍰PD1 (n=6), cDC1v (n=7), cDC1v+ ⍰CTLA4 + ⍰PD1 (n=9), LP: Iso (n=6), LP: ⍰CTLA4 + ⍰PD1 (n=8), LP: cDC1v (n=8), LP: cDC1v+ ⍰CTLA4 + ⍰PD1 (n=8). In **C, D, E, F, G, J, K** and **L** data are presented as mean ± SD. Significance was determined by unpaired t-test in **C**, Mann-Whitney test for comparison of cDC1s and unpaired T-test for all other comparisons in **D**, Mann-Whitney test for comparison of CD4^+^ T cells in **E** and unpaired-T test for all other comparisons in **E**, unpaired T-test in **F** and **G**, log-rank test in **B** and **H**, two-way ANOVA with Tukey’s test in **J**, one-way ANOVA with Sidak’s multiple comparisons test for comparison of mice with pancreatitis and Kruskal-Wallis test for all other comparisons in **K**, by Kruskal-Wallis with Dunn’s multiple comparisons test in **L**. * P <0.05, ** P <0.01, *** P <0.001, **** P <0.0001, *ns*: not significant. Scale bars indicate 100 μm. See related supplementary figures S13-S16. Note: Isotype controls used in Figure 5B, 5H, 5I and 5J are repeated from Fig. 4E, 4I, 4J and 4K respectively to allow for direct comparisons.

The iKPC* mice treated with cDC1 vaccine (iKPC*-cell lysate: AgS) coupled with αCTLA4+αPD1 iCBT exhibited lower tumor weights compared to the isotype treated mice **(Figure 5C)** associated with an increase in the cDC1s and a concomitant decrease in cDC2s frequencies compared to the isotype treated mice **(Figure 5D)**. The cDC1 vaccination with αCTLA4+αPD1 iCBT demonstrated a higher proportion of CD8^+^ T cells, with expression of proliferation marker (Ki67), granzyme B and T-bet, accompanied by a decrease in CD4^+^Foxp3^+^ Tregs **(Figure 5E-5F)**. Although no difference in the Th_1_ population was observed, the cDC1 vaccine with αCTLA4+αPD1 iCBT showed an increase in the CD8/Treg ratio and increasing trend of Th_1_/Treg ratio **(Figure 5G)**. Analysis of exhaustion markers revealed a decrease in frequencies of CD4^+^PD1^+^ and CD8^+^PD1^+^ T cells, whereas insignificant differences in TIM3 expression was observed **(Supplementary Figure S14F and S14G)**. Analysis of CD69, which is associated with memory and activated T cells (*31, 32*), revealed an increase in frequencies of CD3^+^CD69^+^ and CD8^+^CD69^+^ T cells, likely contributing to anti-tumor response and preventing tumor relapse **(Supplementary Figure S14H)**. Collectively, our results indicate that PDAC is likely resistant to iCBT due to the lack of antigen-presenting cDC1s in the TME. cDC1 vaccination rendered PDAC sensitive to iCBT with PDAC eradication observed in some of the mice.

To evaluate the curative potential of cDC1 vaccine, we next conducted a long-term survival study with multiple doses of the cDC1 vaccine in a different PDAC mouse model, which involved orthotopic implantation of a KPC tumor-derived cell line (KPC-689) (**Figure 5A**). Six to eight weeks-old C57BL/6J mice were orthotopically injected with KPC-689 GFP-Luc cells. These mice were subsequently subjected to CD103^+^ cDC1 vaccine (KPC-689 cell lysate AgS, 4 doses), and treated with αCTLA4+αPD1 iCBT (**Figure 5A**). In the KPC-689 PDAC mice, cDC1 vaccine+ iCBT and LP+ cDC1 vaccine+ iCBT led to clearance of PDAC (**Figure 5H-5L and Supplementary Figure S15A-15F**). Next, we probed whether such treatment regimen induced a functional T cell memory. We re-challenged the cured mice on day 85 after regression of PDAC with KPC-689 implantation in the pancreas **(Figure 5H-5J).** IVIS imaging and autopsy after 114 days after tumor re-challenge did not reveal emergence of PDAC (**Figure 5I-5J, Supplementary Figure S16A**). Administration of cDC1 vaccine by itself also increased overall survival of KPC-689 mice with and without LP (**Figure 5H and Supplementary Figure S15B-S15C**). Both KPC-689 tumor cell lysates with and without GFP-Luc demonstrated similar levels of CD8^+^ T cell activation, indicating that GFP does not likely contribute to enhanced tumor clearance **(Supplementary Figure S15F and S16B-S16E**).

Next, we evaluated the cDC1 vaccine (KPC-689 parental cell line lysate: AgS) and αCTLA4+αPD1 iCBT on the autochthonous KPC mice starting at 12w after birth following the establishment of PDAC (**Figure 6A**). cDC1 vaccine with αCTLA4+αPD1 iCBT suppressed PDAC progression and improved the median survival of KPC mice by 15 weeks (**Figure 6B-6C**). Analysis of tumor growth by MRI and age matched analysis by histology demonstrated significant PDAC suppression, accompanied by an increase in CD3^+^, CD4^+^ and CD8^+^ T cells in the pancreas of cDC1v + αCTLA4+ αPD1 iCBT treated KPC mice (**Figure 6D-6H, Supplementary Figure S17A and S17B**). Further, 29% of the mice treated with cDC1 vaccine and αCTLA4+ αPD1 iCBT demonstrated long term survival up to 250 days with minimal evidence of PDAC with greater than 50% normal pancreatic tissue in these mice **(Supplementary Figure S17A-S17B)**.

**Figure 6:**
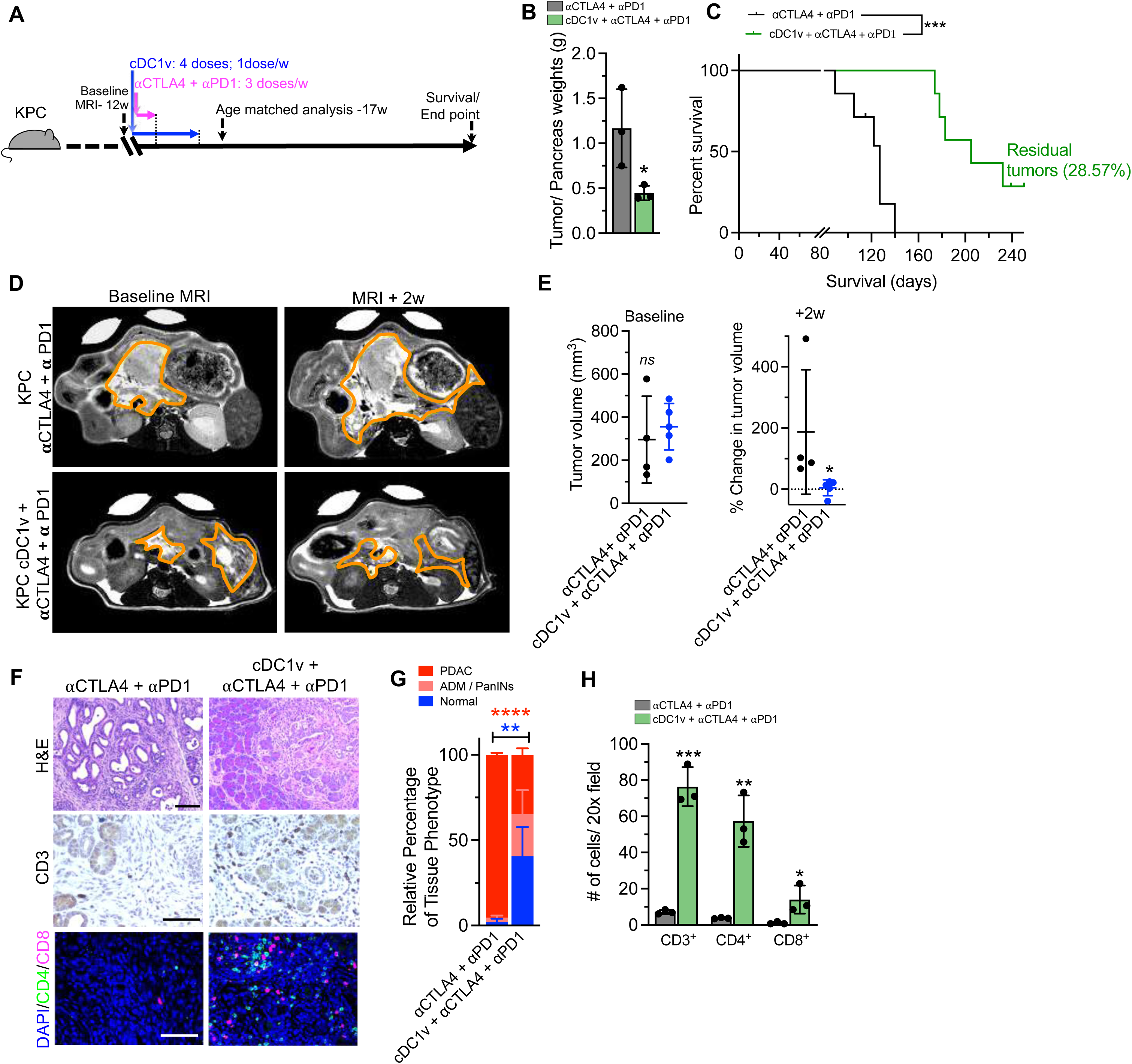
cDC1 vaccine sensitizes KPC mice to checkpoint blockade immunotherapy. (**A**) Schematic representation of treatment of KPC mice with cDC1 vaccine (KPC-689 parental cell lysate used as AgS) and ⍰CTLA4 + ⍰PD1. (**B**) Tumor/ pancreas weights of KPC mice with indicated treatments sacrificed at 17 weeks (n=3/ group). (**C**) Kaplan-Meier survival curve of indicated groups. KPC tumors treated with ⍰CTLA4 + ⍰PD1 (n=7) and cDC1v+ ⍰CTLA4 + ⍰PD1 (n=7). (**D**) Baseline (12 w) (left panel) and post ⍰CTLA4 + ⍰PD1 (n=4) or cDC1v + ⍰CTLA4 + ⍰PD1 (n=5) treatment (2w after treatment) (right panel) MRI imaging of tumors. **(E)** Baseline tumor volumes (left panel) and change in tumor volumes (right panel) of ⍰CTLA4 + ⍰PD1 (n=4) or cDC1v + ⍰CTLA4 + ⍰PD1 (n=5) treated mice. (**F-H**) Age-matched analysis (at 17w) of tumor/ pancreas tissue from ⍰CTLA4 + ⍰PD1 and cDC1v + ⍰CTLA4 + ⍰PD1 treated KPC mice (n=3/ group). Representative H&E, CD3 immunostaining and CD4, CD8 and DAPI immunostaining (**F**), with quantification of H&E (**G**) and immunostaining (**H**). In **B, E, G** and **H** data are presented as mean ± SD. Significance was determined by unpaired T-test in **B** and **H**, unpaired T-test for comparison of baseline tumor volumes and Mann-Whitney test for analysis of change in tumor volumes in **E,** two-way ANOVA with Tukey’s test in **G**, log-rank test in **C**. * P <0.05, ** P <0.01, *** P <0.001, **** P <0.0001, *ns*: not significant. Scale bars indicate 100 μm. See related supplementary figure S17.

### CD103^+^ cDC1 vaccination and immune checkpoint blockade therapy shapes TCR repertoire of tumor infiltrating CD8^+^ T cells in PDAC

To analyze the T-cell receptor (TCR) repertoire associated with anti-tumor CD8 response following cDC1 vaccine and αCTLA4+αPD1 iCBT treatment, we performed high throughput sequencing of the TCRβ CDR3 region of tumor infiltrating CD8^+^ T cells in orthotopic iKPC* tumor bearing mice **(Figure 7A and 7B)**. One week following treatment with cDC1 vaccine and αCTLA4+αPD1 iCBT treatment, the cDC1 vaccine treatment alone did not result in alteration of CD8 frequencies whereas combination with iCBT resulted in significant increase in frequency of intra-tumoral CD8^+^ T cell infiltrates **(Figure 7C)**. cDC1v (loaded with iKPC* PDAC cell lysate) + αCTLA4 + αPD1 iCBT resulted in an increase in CDR3 diversity of tumor infiltrating CD8^+^ T cells **(Figure 7D)**, indicating successful activation of oligoclonal tumor-directed T cell clones. For an in-depth analysis of the distribution, ranking and group average for each clone rank was calculated. A cumulative frequency of 50% was reached in a much larger number of unique clones in the cDC1v + αCTLA4+ αPD1 iCBT associated CD8^+^ T cells compared to the isotype and the cDC1v treatment groups **(Figure 7E-F)**. Our data suggests that cDC1 vaccine did not change the T cell expansion in tumor, but rather their TCR repertoire (priming) **(Figure 7G),** probably due to the immunosuppressive TME, whereas combination of αCTLA4 + αPD1 iCBT potentially broadens it reflecting successful priming in lymphoid tissues, increased migration and subsequent reinvigoration of T cells with unique oligoclonal TCR repertoire in the tumor following iCBT(*33-35*) **(Figure 7E-G).**

**Figure 7:**
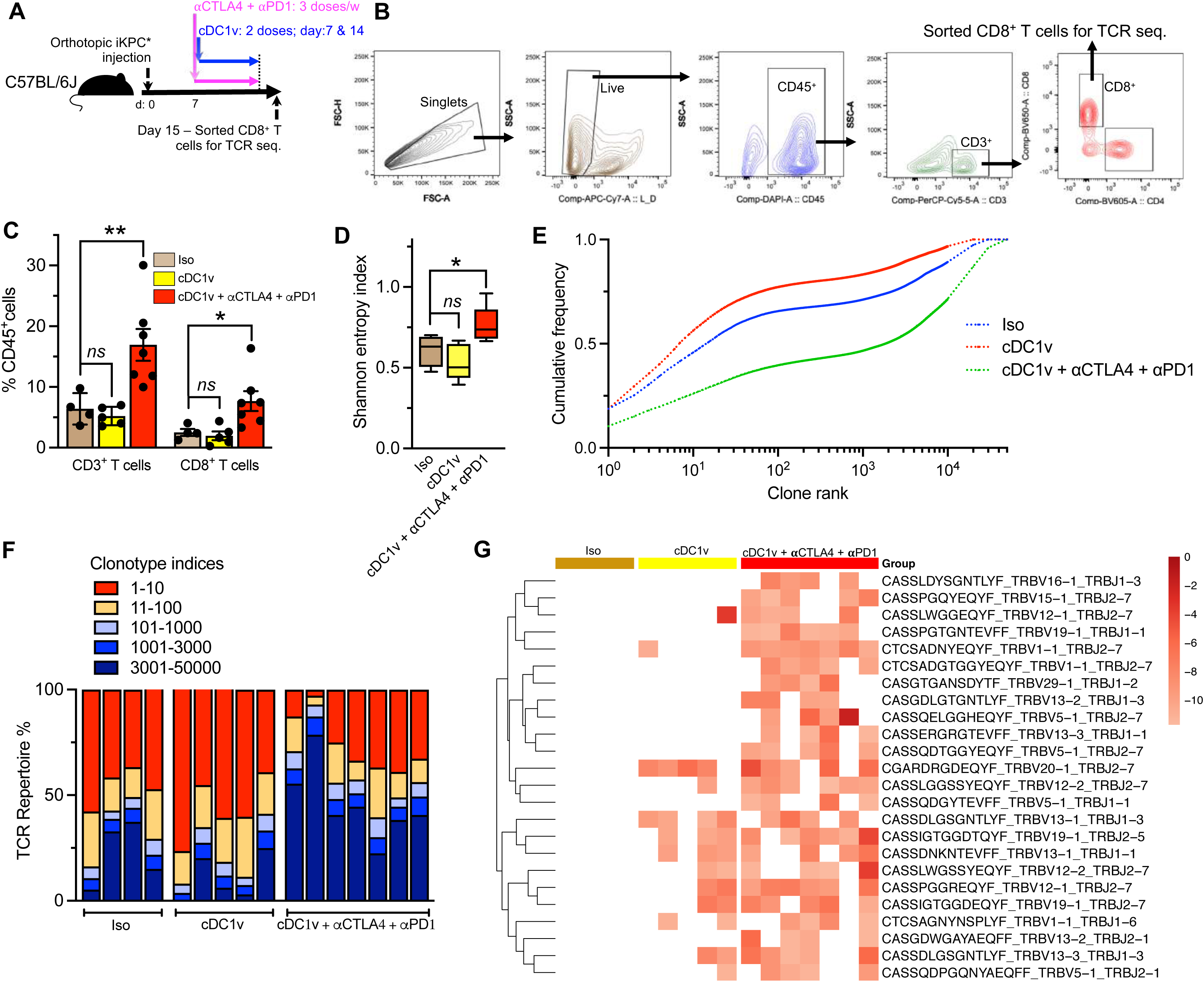
CD103^+^ cDC1 vaccination and immune checkpoint blockade therapy shapes TCR repertoire of tumor infiltrating CD8^+^ T cells in PDAC. (**A**) Schematic representation of orthotopic injection of iKPC* cell line in C57BL/6J mice treated with Isotype (n=4), cDC1 vaccine (n=5), cDC1v + ⍰CTLA4 + ⍰PD1 (n=7). TCR sequencing analysis of tumor infiltrating CD8^+^ T cells was performed on day 15. (**B**) Gating strategy for sorting CD8^+^ T cells for TCR sequencing analysis. (**C**) Flow cytometry analysis of indicated groups. CD3^+^ (T cells); CD3^+^, CD8^+^ (CD8^+^ T cells) were measured as a percentage of CD45^+^ cells. (**D**) Clonality was determined by Shannon entropy index in indicated groups. (**E**) Average frequency distributions for each group were computed and CD8^+^ T cell clones were ranked according to frequency for each mouse. The cumulative frequency was plotted with each clone rank. (**F**) Distribution of the most abundant clonotypes by sample. (**G**) Hierarchical clustering was performed to identify unique CDR3 AA sequence clones present in >50% of animals in the cDC1v + ⍰CTLA4 + ⍰PD1 group. In **C**, data are presented as mean ± SD, as box and whisker plot (min-max) in **D**, as mean in **E** and **F**. Significance was determined by one-way ANOVA with Dunnett’s multiple comparisons test in **C** and **D**. * P <0.05, ** P <0.01, *ns*: not significant. See related supplementary figure S18.

Consistent with increase in TCR diversity, cDC1v + αCTLA4 + αPD1 iCBT treated tumors had expansion of their most abundant clonetypes **(Figure 7F and Supplementary Figure S18A)**. We next analyzed the CDR3 region length distribution of the different treatment groups. Using Gaussian fit for all the treatment groups, the null hypothesis that “One curve fits all datasets” was rejected with a P<0.0001 **(Supplementary Figure S18B)**. The cDC1v + αCTLA4 + αPD1 iCBT treatment demonstrated increased CDR3 sequence counts of 12-16 amino acid (AA) lengths **(Supplementary Figure S18B)**. Next, to identify whether unique CDR3 sequences enriched in response to cDC1v + αCTLA4 + αPD1 iCBT treatment, we performed hierarchical clustering and picked clones that were enriched in atleast 50% of the treated mice, within the top 100 clones with the combination therapy **(Figure 6G and Supplementary Figure S18C)**. We identified 24 unique CDR3 sequences enriched in the mice that received cDC1v + αCTLA4 + αPD1 iCBT treatment. Among them, CTCSADNYEQYF (TRBV1-1_TRBJ2-7) sequence was enriched in 100% of the mice that received the combination **(Supplementary Figure S18C)**.

### Tumor infiltrating CD11c^+^ dendritic cells correlate with CD8^+^ T cell accumulation and increased overall survival of PDAC patients

Based on the impact of DCs in modulating T cell response and survival in murine PDAC, we next assessed the contribution of DCs in the setting of human PDAC. We preformed immunostaining for CD4, CD8, Foxp3 and CD11c in 120 treatment naïve patients with PDAC. Evaluation of CD4^+^, CD8^+^, Tregs and CD11c^+^ cells demonstrated a significant correlation (R^2^ = 0.66) between tumor infiltrating CD8^+^ T cells and CD11c^+^ dendritic cells (**Figure 8A and 8B**). Presence of DCs had a weaker correlation with CD4^+^ T cells (R^2^ = 0.51) and Tregs (R^2^ = 0.2) (**Figure 8A and 8B**). We next divided tumors into ‘high’ and ‘low’ level categories based on the median (**Supplementary Figure S19A**). CD11c ‘high’ patients had a significantly longer disease specific survival (DSS) compared to CD11c ‘low’ PDAC patients (**Figure 8C**). Although there was a good correlation between tumor infiltrating CD11c^+^ DCs and CD8^+^ T cells, the number of CD8^+^ T cells by itself did not predict survival of PDAC patients (**Figure 8D**), indicating that the quality (specificity + affinity) of CTLs rather than their numbers is integral (priming of tumor-directed CTLs). Similarly, CD4^+^ T cells and Tregs infiltration did not predict survival in human PDAC tumors (**Supplementary Figure S19B and S19C**). Next, we analyzed the DC infiltration in 173 treatment naïve patients (TCGA-PanCanAtlas dataset). While the gene encoding the generic DC marker, CD11c-integrin subunit alpha X (*Itgax*) did not predict higher survival of PDAC patients in the TCGA dataset **(Supplementary Figure S19D)**, basic leucine zipper ATF-like transcription factor 3 (*Batf3*) expression, a specific marker for human antigen presenting cDC1s, predicted increased survival of PDAC patients **(Figure 8E)**.

**Figure 8:**
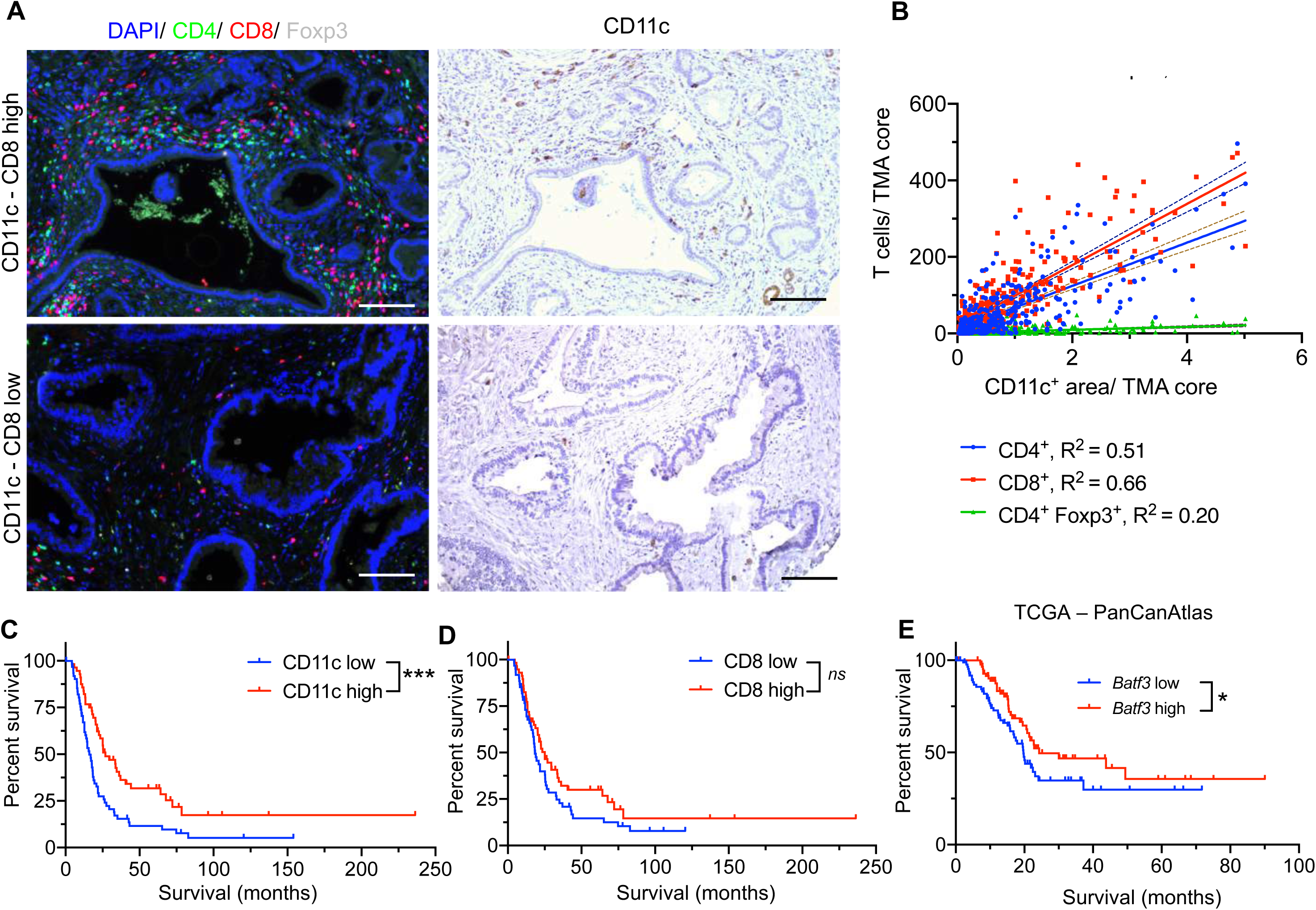
Tumor infiltrating dendritic cells correlate with CD8^+^ T cell infiltration and better prognosis in human PDAC. (**A**) Representative CD4^+^, CD8^+^, Foxp3^+^ and CD11c^+^ immunostaining in human PDAC TMA samples. (**B**) Linear regression analysis of CD11c^+^ DCs with each T cell group: CD4^+^ T cells (blue), CD8^+^ T (red) and CD4^+^, Foxp3^+^ T cells (green) in each tissue sample; n=120 PDAC patients (316 tissue sample cores). (**C-D**) Kaplan-Meier survival curves for disease-specific survival (DSS) of CD11c high (n=56) and CD11c low (n=63) tumors (**C**), and CD8 high (n=59) and CD8 low (n=61) tumors (**D**). (**E**) Kaplan-Meier survival curves for DSS of PDAC patients with *Batf3* high (n=88) vs. *Batf3* low (n=89) expression based on RNA-seq. analysis of samples from TCGA – PanCanAtlas dataset. Significance was determined by Log-rank test in **C, D** and **E.** R^2^ represents square of correlation co-efficient from regression analysis in **B**. Scale bars indicate 100 μm. * P <0.05, *** P <0.001, *ns*: not significant. See related supplementary figure S19.

## Discussion

Pancreatic cancer is associated with compromised tumor immunity with dysfunctional T cells, as also demonstrated in this study (*36*) (*17*). We show that when pancreatic cancer is associated with tissue injury and inflammation due to pancreatitis, the immune profile is altered with increased number of activated CD11c^+^ DCs and cDC1s. In WT mice with pancreatitis, mechanisms of peripheral tolerance likely result in T cell suppression in response to antigen presenting cDC1s that present self-antigens resulting from auto-digestive process via activation of proteolytic enzymes (*25, 29*). We show that this feature is inherent to the tissue injury and inflammation induced by pancreatitis in WT mice as part of the immune response to protect functional parenchyma from autoimmune damage. Interestingly, in the absence of pancreatitis, the majority of DCs were present within the lymphoid structures but not within PanINs or PDAC. Pancreatitis associated with PanINs and PDAC presumably resulted in migration of activated DCs into PanINs and the intra-tumoral space of PDAC.

In the context of pancreatic cancer, the activated CD11c^+^ DCs that emerge due to pancreatitis are primed to present tumor antigens but failed to elicit an anti-tumor CD8^+^ T cell response due to inhibitory CD4^+^ T cells. Lack of CD4^+^ cells, or relieving the suppressive effects of Tregs via anti-CTLA-4 and/or anti-PD-1 iCBT enabled effector CD8^+^ T cells and Th_1_ response which led to dramatic suppression of PDAC with long-lasting cures in different mouse models. Collectively, this study demonstrates that the immuno-suppressive nature of pancreatic cancer is likely due to lack of proper priming by cDC1s.

As reported in this study and by others (*17*), depletion of CD4^+^ or CD8^+^ T cells in PDAC was shown not to impact PDAC progression in mouse models. Further, other studies that used pancreatitis-accelerated pancreatic cancer mouse models as equivalent surrogates of spontaneous GEMMs conclude that CD4^+^ T cells and Th_17_ cells play a PDAC-promoting and Tregs play a PDAC-inhibitory role (*21-23*). The potential role of overlaying pancreatitis in altering the immune microenvironment to impact progression of pancreatic cancer was not explored in these studies. Our work here clarifies this issue, and illustrates that IME of pancreatic cancer with overlaying pancreatitis is different from pancreatic cancer without pancreatitis. The immune microenvironment of pancreatitis mechanistically highlights an opportunity to realize the efficacy of iCBT in pancreatic cancer.

Monocyte derived DC (Mo-DC) vaccines have been used in pre-clinical models (*37, 38*) and clinical trials for different cancers with limited efficacy (*39-41*).The Mo-DCs are derived from ex-vivo cultures of immune cells isolated from peripheral blood upon treatment with GM-CSF and IL-4, and subsequently stimulated with tumor antigens. Despite demonstrating ability to cross-prime T cells, Mo-DC vaccine may exhibit suboptimal vaccine efficacy due to limited migratory capacity to lymph nodes (*42*).

Tumor infiltrating cDC1s play a critical role in CD8^+^ T cell priming and mediating anti-tumor immune responses, and cDC2s are implicated in priming T helper cells and mediating Th_2_ responses (*28, 43, 44*). Efficacy of cDC1 vaccine has been described in osteosarcoma and melanoma (*37*). Here we report that cDC1 vaccine in combination with αCTLA4+αPD1 iCBT eradicates PDAC. Our findings also show that administration of CD103^+^ cDC1 vaccine as monotherapy resulted in marginal disease control by itself and increased the survival of mice with PDAC. iCBT achieved PDAC eradication associated with cytotoxic CD8^+^ T cells and Th_1_ activation, suppression of Tregs, and expansion of CD69^+^ T cells with activated/memory phenotype (*45*). Our findings are further supported by a report which suggested that lack of DCs in the PDAC TME could lead to dysfunctional immune surveillance (*28*).

Lack of tumor growth in mice after re-challenge, upon cDC1 vaccine + αCTLA4 +αPD1 iCBT following eradication of PDAC, illustrated potential contribution of functional T cell memory. Analysis of the tumor infiltrating CD8 clones indicate that cDC1 vaccine + αCTLA4 + αPD1 iCBT broadens the TCR repertoire and resulted in an increase in unique CD8^+^ T cell clones infiltrating PDAC. We identified unique CDR3 AA sequences in tumor infiltrating CD8^+^ T cells that could potentially inform future T cell-transfer therapies. It is important to note that we observed an augmented oligoclonal CD8^+^ T cells response with unique CDR3 sequences, which likely represents the fact we used total tumors/cancer cells lysate to induce presentation of multiple antigens by activated cDC1s, versus using a single tumor specific peptide (*33, 34*). Use of a single peptide to generate antigen presenting cDC1s would likely generate more restricted clonal response, as demonstrated in another study (*46*).

In conclusion, our study provides a comprehensive and context dependent analysis of T cells and its interaction with DCs in the PDAC IME. The presence of antigen presenting cDC1s recruited either by pancreatitis or by exogenous administration of CD103^+^ cDC1 vaccine, sensitizes the PDAC TME to iCBT. Our study suggests that patients with a history of pancreatitis or concurrent pancreatitis may benefit from iCBT, and also provides rationale for combining cDC1 vaccination with iCBT in PDAC. Further, our study provides encouraging pre-clinical data for the development of human cDC1 vaccine and their combination with iCBT for the treatment of PDAC.

## Material and methods

### Animal studies

The genotyping and tumor progression of the *Pdx1-cre; LSL-Kras^G12D/+^; Trp53^R172H/+^* (KPC) and *Pdx1-cre; LSL-Kras^G12D/+^* (KC) mice have been described previously (*47*). We crossed KPC to CD4^-/-^ (*Cd4^tm1Mak^*)(*48*) or CD8^-/-^ (*Cd8^tm1Mak^*)(*49*) mice purchased from Jackson Laboratory to obtain KC, KC CD4^-/-^, KC CD8^-/-^, KPC, KPC CD4^-/-^ and KPC CD8^-/-^ mice. For orthotopic experiments, 6-8 week old C57BL/6J mice were injected with 5 x 10^5^ primary PDAC cell lines in 20μL PBS, either KPC-689 GFP-Luc cell line (*50*) or iKPC* cell line (*51*) (kindly provided by Dr. Haoqiang Ying) in the tail of the pancreas. The iKPC* cell line has a tetracycline-inducible *tetO-LSL Kras^G12D/+^* allele and implanted mice were maintained on doxycycline (Dox) water (Dox 2 g/L, sucrose 20 g/L), initiated with orthotopic injection and maintained throughout the experiment. For the tumor re-challenge study, 5 x 10^5^ KPC-689 GFP-Luc cells were injected into the pancreas at day 85 following the initial injection. Tumor radiance (photons s^−1^ cm^−2^ sr^−1^) was monitored for the KPC-689 GFP-Luc cell line injection using IVIS imaging (Xenogen spectrum) under uniform conditions across all experimental groups. Mice were injected with D-luciferin (Gold Biotechnology, LUCK-100) (100 mg/kg, at 10 mg/ml concentration) intraperitoneally and imaged under isoflurane anesthesia 10 minutes following injection. In the autochthonous KPC mice, MRI measurements of tumor volumes were determined before treatment start and subsequently followed up after 2 weeks to monitor tumor growth using Bruker 7T MRI. For pancreatitis induction, caerulein was injected at a final volume of 100uL (dose – 50 μg/kg per mouse), four times a day (every 6 hours), on alternate days for short-term pancreatitis and three times a week for 3 weeks to induce long-term pancreatitis as described previously (*22*). For experiments to neutralize CD11c^+^ DCs, 500 μg of anti-mouse CD11c (ThermoFisher scientific, clone N418, CUST05168) (1 mg/mL concentration) was injected intraperitoneally on days 1, 5, 10 and 15. Isotype controls for anti-CD11c treated mice were injected with Armenian hamster IgG (BioXcell, BE0260) in the same route, time and dosing as the neutralizing antibody. Depletion of CD8^+^ T cells were performed simultaneously starting at the time of SP induction by administration of 200µg of anti-mouse CD8 (BioXcell, 53-6.7, BE0004-1) twice per week in 100 µL of PBS intraperitoneally for the entire course of the experiments. For checkpoint immunotherapy, anti-mouse CTLA4 (BioXcell, clone 9H10, BE0131) and/ or anti-mouse PD-1 (BioXcell, clone 29F.1A12, BE0273) were injected three times intra-peritoneally as indicated, starting dose 200 μg followed by 2 doses of 100 μg each (1 mg/mL concentration). Isotype controls for anti-CTLA4 and anti-PD1 treated mice were injected with a combination of Rat IgG2a (BioXcell, clone 2A3, BE0089) and Syrian hamster IgG (BioXcell, BE0087) in the same route, time and dosing as the neutralizing antibodies. For DC vaccine, 1.5 – 2 x 10^6^ cDC1s were injected intraperitoneally once every week for 4 weeks in the KPC GEMMs and KPC-689 GFP Luc orthotopic tumor bearing mice and for 2 weeks in the iKPC* orthotopic tumor bearing mice. For TCR sequencing analysis of CD8^+.^T cells, CD8^+^ T cells were sorted from the tumor infiltrating lymphocytes 2 weeks after orthotopic injection of iKPC* cells.

### Cell culture

Primary PDAC cells viz. KPC-689 GFP-Luc (*50*) and iKPC* cells (*51*) have been described previously. KPC-689 GFP-Luc cells were cultured in 10% FBS in RPMI (Corning, 10-040-CV) with 1% penicillin-streptomycin (PS) (Corning, 30-002-CI) and iKPC* cells were cultured in 10% tet-free FBS (Gemini Bio-Products, 100-800) in DMEM (Corning, 10-017-CV) with 1% PS and 1 μg/mL doxycycline (StemCell Technologies 72742). Prior to orthotopic injection of primary PDAC cell lines, cells were confirmed to be negative for mycoplasma using LookOut® Mycoplasma PCR Detection Kit (Sigma-Aldrich, MP0035).

### cDC1 vaccine

Preparation of the CD103^+^ cDC1 vaccine has been described previously (*37, 52*). Briefly, 6-10 weeks old C57BL/6J murine bone marrow culture was established following RBC lysis at a concentration of 1.5 x 10^6^ cells/mL in cRPMI (10% heat-inactivated fetal bovine serum (FBS) (Atlanta Biologicals, Atlanta, Georgia, USA), 1% penicillin-streptomycin (PS), 1 mM sodium pyruvate, and 50 µM β-mercaptoethanol) supplemented with 50ng/mL hFIt3-L (PeproTech, 10773-618) and 2ng/mL GM-CSF (PeproTech, 315-03). The culture was supplemented on day 5 with a 50% increase in cRPMI by volume and subsequently, the non-adherent cells are re-plated at a concentration of 3 x 10^5^ cells/mL supplemented with the same amount of hFIt3-L and GM-CSF on day 9. The supernatant with non-adherent cells was collected on day 15-17 for co-stimulation with tumor lysate. The cell pellet was stained for the following markers: CD11c, B220, CD24, CD172a and CD103. The cDC1s (live/dead stain^-^, CD11c^+^, CD24^+^, CD172^-^, CD103^+^, B220^-^) were sorted on FACS Aria Fusion sorter and plated at 2-4 x 10^6^ cells/mL in cRPMI, incubated with lysate prepared from the antigen source (KPC-689 GFP-Luc cells, iKPC* cells, WT-LP pancreas lysate or iKPC*-LP tumor lysate in a Lysate: cDC1 ratio of 2:1). For preparation of the lysate, cell pellets were freeze-thawed three times rapidly from -80°C to 37°C. Subsequently, ultrasonic disruption of cells was performed using Heat Systems-Ultrasonics W-385 Sonicator Ultrasonic Processor and Probe set to 40% duty cycle with a microtip limit of 5. The probe was inserted into the sample and was ultrasonicated for 20 s with one pulse increments. For generation of lysates from WT pancreas and iKPC* orthotopic tumors with LP, tumors/pancreas were minced and digested in 1.5 mg/mL Collagenase P (Sigma-Aldrich) in 5 mL HBSS at 37°C for 20 minutes. Subsequently, multiple washes were performed in 10% FBS in RPMI with PS and filtered using 70 μm strainer (Corning 352350) and centrifuged at 600g for 3 minutes in room temperature. Lysates were prepared after freeze thaw cycles and ultrasonic disruption as described above. The cDC1s were stimulated with AgS lysates in media supplemented with 20 µg/mL poly I:C (Sigma-Aldrich, P4929) and 20 ng/mL GM-CSF for 4 hours. Cells were washed and resuspended in PBS. Subsequently, 1.5 - 2 x 10^6^ cDC1s were resuspended in 100 µL PBS and injected intraperitoneally as indicated.

For in-vitro experiments to compare the effects of KPC-689 GFP-Luc tumor lysate vs. KPC-689 parental tumor lysates on T cells, we stimulated cDC1s with poly I:C, the respective tumor lysates (ratio of tumor cell lysate: cDC1s = 2:1) or with both poly I:C and tumor lysates. We plated the T cells and cDC1s at a concentration of 1 x 10^5^ cells/mL, 100µL of each in a 96 well plate. After incubation, the cells were centrifuged at 600g for 3 minutes in room temperature and washed in FACS buffer. Then, the cells are stained with a cocktail of antibodies for CD3, CD8, PD1, CD25 and CFSE for 30 minutes on ice. Cells were fixed in 1.6% formaldehyde and analyzed by flow cytometry. All antibodies were used at a concentration of 1:200 for cDC1 sorting and CD8^+^ T cell stimulation experiments. See **Supplementary Table 1** for antibody information.

### Immunostaining

For single stained immunohistochemistry (IHC), 5 µm thick formalin fixed paraffin embedded (FFPE) slides were deparaffinized and antigen retrieval was performed in indicated buffers at 95°C for 20 minutes. For CK19, CD11b, CD68 and CD11c staining, citrate buffer (pH = 6) was used, whereas for CD4, CD8 and Ki67 staining, Tris-EDTA buffer (pH = 9) was used for antigen retrieval. Subsequently, slides were blocked in 1.5% bovine serum albumin in PBST (0.1% Tween 20) for 30 minutes. Slides were then incubated in 3% H_2_O_2_ in PBS for 15 minutes. Primary antibodies CK19 (Abcam, ab52625, 1:250), CD11b (Abcam, ab13357; 1:500), anti-mouse CD11c (Cell Signaling Technology, CST 97585S, 1:350), anti-human CD11c (Abcam, ab52632, 1:100), anti-mouse CD4 (Abcam, ab183685, 1:400), anti-mouse CD8 (Cell Signaling Technology, 98941s, 1:250) and Ki67 (ThermoScientific, RM-9106-S, 1:100) were diluted in 1.5% BSA in PBST and incubated overnight at 4°C. For all IHC, sections were incubated with biotinylated secondary antibody for 30 minutes followed by ABC kit (VECTASTAIN Elite ABC-HRP kit, Vector Laboratories, PK-6100) for 30 minutes. Next, DAB and counterstaining with hematoxylin were performed and DAB positivity was quantified by examining multiple random visual fields under the microscope. For the thymus and spleens of KPC, KPC CD4^-/-^ and KPC CD8^-/-^ mice, 5 μm-thick cryostat OCT μ sections were fixed in acetone at 4°C for 5 minutes, blocked in 1.5% BSA in PBS for 30 minutes, stained with primary antibodies – anti-mouse CD4 (Abcam, ab183685, 1:400) or anti-mouse CD8 (AbD Serotec, MCA1767T, 1:100) in 1.5% BSA in PBS (1 hour at room temperature) and secondary antibodies (Goat anti-rabbit (H+L), Alexa Fluor Plus 488, ThermoFisher, A32731, 1:250 for CD4 staining or Goat anti-rat IgG (H+L), Alexa Fluor 488, 1:250 for CD8 staining) (30 minutes at room temperature). Immunofluorescence staining performed using Tyramide signaling amplification (TSA) was performed as described elsewhere (*20*). The reagents used and the order of antibody staining are listed in **Supplementary Table 2**.

### Flow cytometry

Tumors or pancreas were minced and digested in collagenase P, 1.5 mg/mL (Sigma-Aldrich, 11213873001) in 5 mL HBSS at 37°C for 20 minutes. Subsequently, multiple washes were performed in cRPMI and filtered using 70 μm strainer (Corning, 352350) and centrifuged at 600g for 3 minutes at 4°C. Cells were washed and resuspended in FACS buffer. Subsequently, cells were incubated in RBC lysis buffer (ThermoFischer, 00-4300) for 5 minutes. Cells were stained with 100 μL surface antibody cocktail diluted in FACS buffer, 20% brilliant stain buffer (BD Bioscience, 566349), Live/dead stain (eBioscience, 65-0865-14) and 50 μ mouse CD16/32 (TONBO biosciences, 40-0161) for 30 minutes on ice, protected from light. For intracellular staining, cells were fixed and permeabilized in Foxp3/Transcription Factor Staining Buffer Set (eBioscience, 00-5523-00) and incubated with intracellular antibodies diluted in Fixation/Permeabilization diluent (eBioscience, 00-5223) for 30 minutes. Subsequently, cells were fixed with fixation buffer (BD Bioscience, 554655) and data were acquired using a BD LSR Fortessa-X20 and analyzed with FlowJo v10.7.1. Antibodies panels used for flow cytometry are listed in **Supplementary Table 3**.

### Mass cytometry

Tumors or pancreas were minced, digested, and stained with anti-CD45 as described earlier in the flow cytometry section. Following RBC lysis, CD45^+^ lymphocytes were flow sorted and 1 x 10^6^ cells were used for staining with antibody cocktail (**Supplementary Table 4**) with anti-mouse CD16/32 (TONBO biosciences, 40-0161) for 30 minutes at room temperature in a final volume of 100uL in MaxPAR cell staining buffer (Fluidigm, 201068). Cisplatin (Fluidigm, 201064) viability staining was added at a 5 µM final concentration in MaxPAR PBS (Fluidigm, 201058). The cells were fixed in 1.6% formaldehyde solution (ThermoFisher, 28906) in MaxPAR PBS for 10 minutes in room temperature. Cells were then incubated in Cell-ID Intercalator Ir (Fluidigm, 201192A) prepared in MaxPAR Fix and Perm Buffer to a final concentration of 125 nM overnight at 4°C. The cells were resuspended in MaxPAR water (Fluidigm, 201069) and analyzed with a Fluidigm Helios Mass Cytometer. Mass cytometry data was initially processed and manually gated in FlowJo (BD Biosciences, version 10.7.1). Live CD45^+^ cells of each sample with the same percentage were exported and utilized for the downstream clustering analysis. We conducted downstream analysis using the approaches described in the R (version 4.0.2) package CyTOF workflow (version 1.7.2). Specifically, R package FlowSOM (version 1.20.0) was employed to computationally define the initial cell clusters using the following parameters: CD45, PD-L1, CD40, CD80, CD19, CD11b, Ly-6G, F4/80, Ly-6C, CD3e, PD-1, CD8a, CD4 and CD11c, followed by identification cell metaclusters based on the heat map. Dimensionality reduction analysis was conducted by t-stochastic neighbour embedding (t-SNE) with R package scatter (version 1.16.2). See **Supplementary Table 4** for antibody information.

### T cell receptor sequencing and data analysis

Total genomic DNA from the sorted CD8^+^ T cells in the indicated groups were extracted using QIAamp DNA Micro Kit (Qiagen, 56304), eluted in a final volume of 50µL buffer AE and submitted for amplification and sequencing of the TCRβ region, survey resolution to Adaptive Biotechnologies (ImmunoSEQ platform). The TCR sequencing data was exported from the immnunoSEQ Analyzer 3.0. Immunarch (0.7.0) (doi: 10.5281/zenodo.3367200) was then employed to perform the various TCR repertoire analysis. The TCR clonotype diversity matrix was calculated using the Shannon entropy index. It used clonotype frequencies to assess the diversity of each sample. The clonality of each sample was also measured using Immunarch. We ranked the clonotypes based on their frequencies and calculated the proportions of groups based on different frequency classifications. CDR3 length distribution was quantified based on the amino acids of the CDR3 region.

To identify the treatment-related clusters of clones, we defined the clones using the combination of CDR3 amino acid sequence, V gene and J gene. For clones with different TCR CDR3 nucleotide sequences but have the same amino acid sequences after translation, we added their proportions. We selected the top 100 clones for each sample in the cDC1v + CTLA4 + PD1 group and then picked the clones detected in at least 4 samples (> 50% of the mice) of this group but were not detected in any Iso group. The heatmap of clones was generated using the *pheatmap* (1.0.12) package under the R environment (4.2.1). The frequency was log-transformed (clones not detected were given a log-transformed frequency of 0).

### PDAC-TMA and TCGA dataset analysis

For the PDAC-TMA dataset, 2-3 cores were selected from FFPE tumor blocks of archived PDAC specimens and TMAs with 1mm^2^ core area were generated. Serial sections were used for CD4-CD8-Foxp3 and CD11c staining. 120 treatment naïve samples were stained to analyze immune infiltration in these tumors. The patient characteristics of these samples are tabulated in **Supplementary Table 5**. TCGA survival analyzes were performed using pancreatic adenocarcinoma (PAAD) gene expression data on treatment 177 naïve samples and clinical data downloaded from UCSC Xena (DOI: 10.1038/s41587-020-0546-8). The gene expression was normalized by logarithm 2 in UCSC Xena. We divided the tumor samples into two groups based on the median gene expression and disease-specific survival (DSS) was plotted in Prism to generate Kaplan-Meier survival plots.

### Statistical analysis

Statistical tests were performed using GraphPad Prism 8 and R-studio. To assess the normality of distribution, Shapiro-Wilk test was used. For comparison of means between two groups with continuous variables, unpaired T-test was used for normal distributions and Mann-Whitney test was used for non-normal distributions. For multiple comparisons, one-way ANOVA was used for groups with normal distribution and Kruskal-Wallis test was used for groups with non-normal distribution. Comparison of relative percentage of histological phenotypes were assessed by two-way ANOVA. Log-rank test was used to compare Kaplan-Meier survival curves. P values are defined throughout the manuscript as: * P<0.05, ** P<0.01, *** P<0.001, **** P<0.0001, *ns*: not significant.

## Acknowledgements

We wish to thank Michelle Kirtley for technical support with managing mouse colony and the South Campus Flow Cytometry Core Lab of MD Anderson Cancer Center for technical support with FACS sorting and analysis. We thank the North Campus Flow Cytometry Core Lab of MD Anderson Cancer center for running mass cytometry. We also thank Dr. Florencia McAllister for providing tissue and feedback on data, Visweshwaran Ravikumar for technical help with TCGA datasets, Swetha Anandan for advice with mass cytometry and Dr. Padmanee Sharma for providing us the commercial vendor information for TCR sequencing. The research conducted in R.K. lab for this manuscript was supported by funds from the MD Anderson Cancer Center. The research related to pancreatic cancer in the Kalluri Laboratory is also supported by the NIH grant P01CA117969. NIH grants R01AI109294 and R01AI133822 to S.S.W. supported the generation of cDC1 vaccine. Other support includes, Flow Cytometry and Cellular Imaging Core Facility by NCI P30CA16672.

## Author contributions

Conceptualization: K.K.M. and R.K.; data curation: K.K.M.; formal analysis: K.K.M., B.L., H.S., and A.M.D.; methodology: K.K.M., and A.M.D.,; investigation: K.K.M. and A.M.D; human PDAC sections and approvals: H.W.; CyTOF analysis: B.L., and K.K.M. TCGA - PanCanAtlas dataset RNA sequencing analysis: B.L. and K.K.M; resources: R.K.; supervision: Y.C., H.W., K.M.M., and R.K.; validation: K.K.M., K.M.M., Y.C.; writing – original draft: K.K.M, and R.K.; writing – review & editing: K.K.M., K.M.M., Y.C., A.M.D., S.S., S.W., and R.K.

---

**Supplementary Figure 1:**
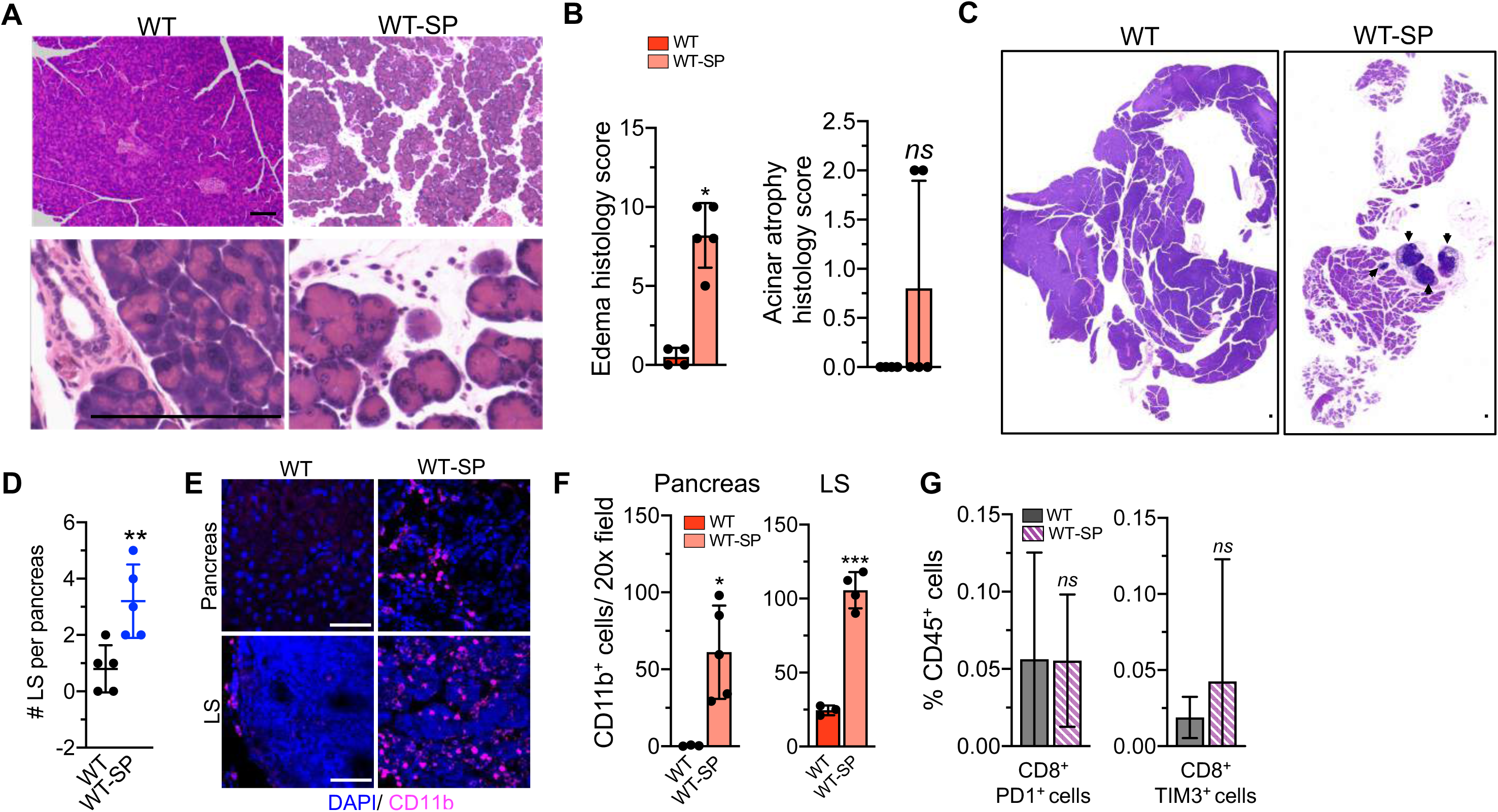
Additional analysis of short-term pancreatitis in wildtype mice. **Related to Figure 1.** (**A-B**) Representative H&E images (**A**), with quantification of tissue damage seen as edema and acinar atrophy (**B**), in WT (n=4) and WT-SP (n=5) mice pancreata. (**C-D**) Representative H&E (**C**), with quantification of number of lymphoid structures (LS) in the pancreas (**D**), of WT and WT-SP mice (n=5/ group). WT (n=20 mice, pancreata from 4 mice were combined per replicate), WT-LP (n=10 mice, pancreata from 2 mice were combined per replicate) mice. **(E-F)** Representative CD11b immunostaining (**E**) with quantification of pancreas and LS in pancreas (**F**) of WT and WT-SP mice (n=3-5/ group). (**G**) Immunophenotyping analysis of T cell populations in WT (n=20) and WT-SP (n=10) mice by flow cytometry. Relative frequencies of immune cells in indicated groups. CD8^+^PD1^+^ and CD8^+^TIM3^+^ cells as a percentage of CD45^+^ cells. In **B, D, F and G**, data are presented as mean ± SD. Significance was determined by Mann-Whitney test in **B** and unpaired t-test in **D, F and G.** *P <0.05, **P <0.01, *** P <0.001 *ns*: not significant. Scale bars indicate 100 μm.

**Supplementary Figure 2:**
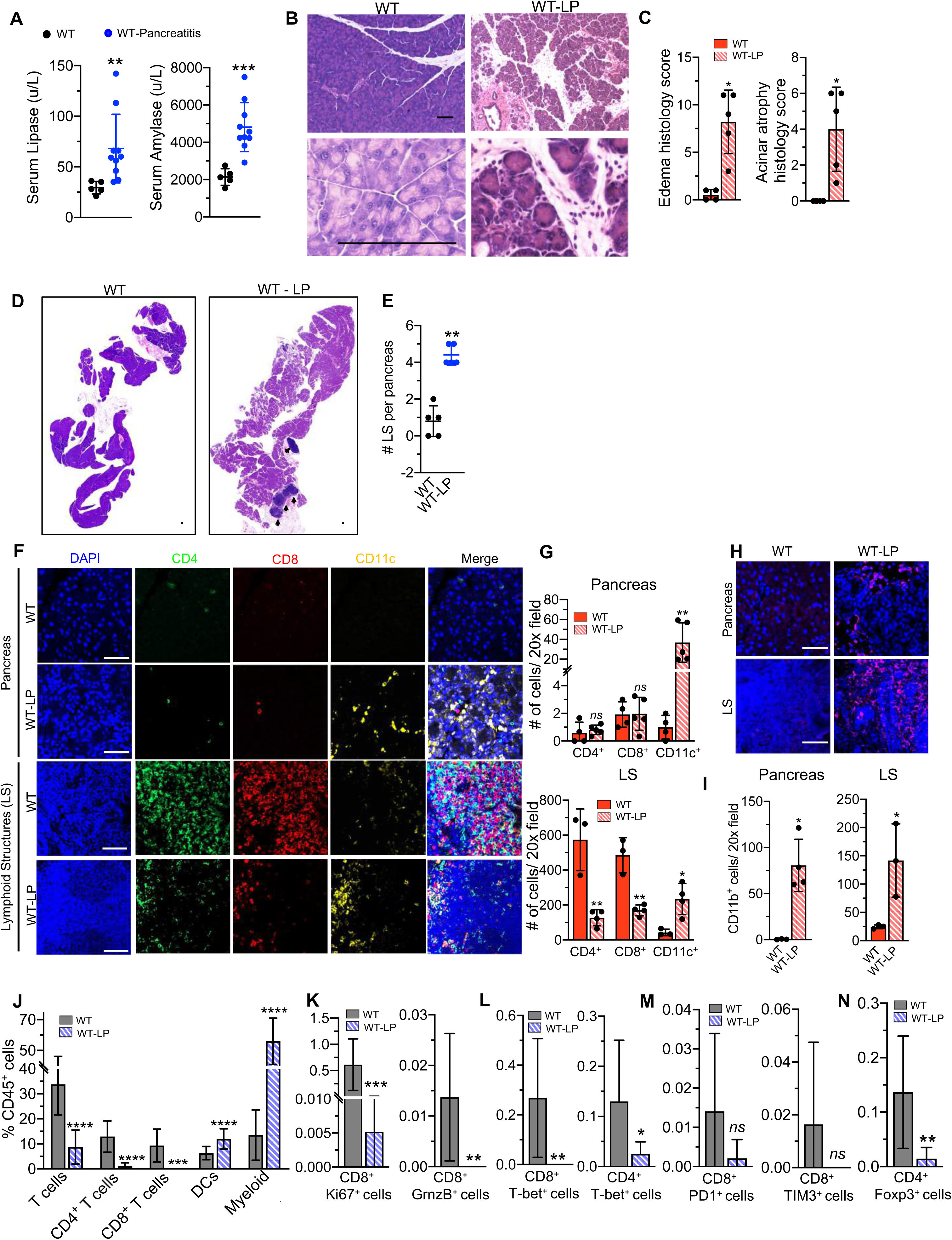
Analysis of pancreatic immune infiltrates of wildtype mice with long-term pancreatitis. **Related to Figure 1.** (**A**) Serum amylase and lipase levels in WT (n=5) and WT-LP (n=10) mice one week following start of caerulein injections. (**B-C**) Representative H&E images (**B**), with quantification of tissue damage seen as edema and acinar atrophy (**C**), in WT (n=4) and WT-LP (n=5) mice pancreas. (**D-E**) Representative H&E (**D**), with quantification of lymphoid structures (LS) in the pancreas (**E**) of WT and WT-LP mice pancreas (n=5/ group). Note: Same WT controls from Supplementary Fig. S1A-C. **(F-G)** Representative CD4, CD8, CD11c immunostaining (**F**), with quantification of pancreas and LS in pancreas (**G**) of WT and WT-LP mice (n=3-5/ mice). **(H-I)** Representative CD11b immunostaining (**H**), with quantification of pancreas and LS in pancreas (**I**) of WT and WT-LP mice (n=3-5/ mice). (**J**) Immune cell populations from pancreas of WT (n=20, pancreas from 4 mice combined per replicate) and WT-LP mice (n=10, pancreas from 2 mice combined per replicate) by flow cytometry. (**K-N**) Immunophenotyping analysis of T cell populations in WT and WT-LP mice by flowcytometry. Relative frequencies of immune cells in indicated groups. CD8^+^, Ki67^+^ and CD8^+^, GrnzB^+^ (**K**); CD8^+^, T-bet^+^ and CD4^+^, T-bet^+^ (**L**); CD8^+^, PD1^+^ and CD8^+^, TIM3^+^ (**M**); and CD4^+^, Foxp3^+^ cells (**N**); measured as a percentage of CD45^+^ cells. Data are presented as mean ± SD in **A, C, E, G, I, J, K, L, M and N**. Significance was determined by Mann-Whitney test for comparison of serum lipase and unpaired T-test for comparison of serum lipase in **A**, Mann-Whitney test in **C and E**, unpaired T-test in **G, I, J, K, L, M and N**. *P <0.05, **P <0.01, *** P <0.001, ****P <0.0001, *ns*: not significant. Scale bars indicate 100 μm. Note that same WT mice pancreas tissue from Figure 1 and supplementary figure S1 are used to allow direct histological and immunostaining analysis.

**Supplementary Figure 3:**
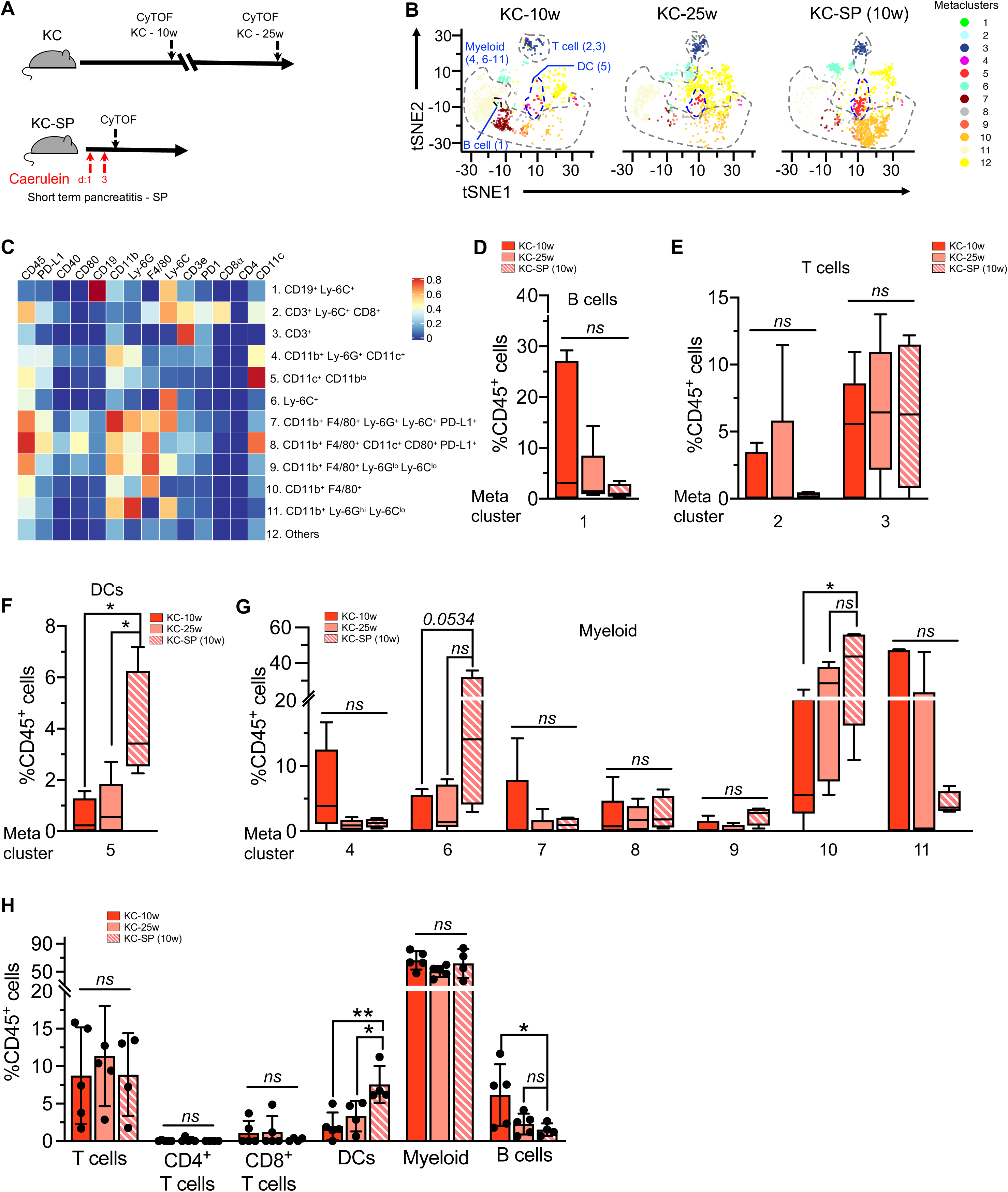
CyTOF analysis of KC mice with short-term pancreatitis. **Related to Figure 2.** (**A**) Schematic representation of induction of short-term pancreatitis (SP) and CyTOF analysis (each red arrow indicates injection of caerulein 4 times per day, 6 hours apart) in KC mice with age and stage matched controls. (**B**) Representative viSNE plots on CD45^+^ cells in KC-10w (n=5), KC-25w (n=5) and KC-SP (10w) (n=4) mice pancreata assessed by CyTOF. (**C**) Heat map of KC mice pancreata infiltrating immune cell metaclusters (MCs) displaying expression values of individual parameters normalized to the maximum mean value across MCs. (**D-G**) Relative frequencies of B cell MCs (**D**), T cell MCs (**E**), dendritic cell MCs (**F**), and myeloid MCs (**G**) by unsupervised clustering of CyTOF markers in KC-10w (n=5), KC-25w (n=5) and KC-SP (10w) (n=4) mice pancreata. (**H**) Immune cell populations identified by Boolean gating of CyTOF markers in indicated groups. In **D, E, F** and **G** data are presented as box and whisker plots (min to max) and as mean ± SD in **H**. Significance was determined by Kruskal-Wallis with Dunn’s multiple comparisons test in **D**, Kruskal-Wallis with Dunn’s multiple comparisons test for MC 2 and one-way ANOVA with Dunnett’s multiple comparisons test for MC 3 in **E**, one-way ANOVA with Dunnett’s multiple comparisons test for MC 5 in **F**, one-way ANOVA with Dunnett’s multiple comparisons test for MCs 4, 10 and Kruskal-Wallis with Dunn’s multiple comparisons test for MCs 6, 7, 8, 9 in **G**, one-way ANOVA with Dunnett’s multiple comparisons test for T cells, DCs, myeloid, B cells comparisons and Kruskal-Wallis with Dunn’s multiple comparisons test for CD4^+^ T cells, CD8^+^ T cells comparisons in **H**. *P <0.05,**P<0.01, *ns*: not significant.

**Supplementary Figure 4:**
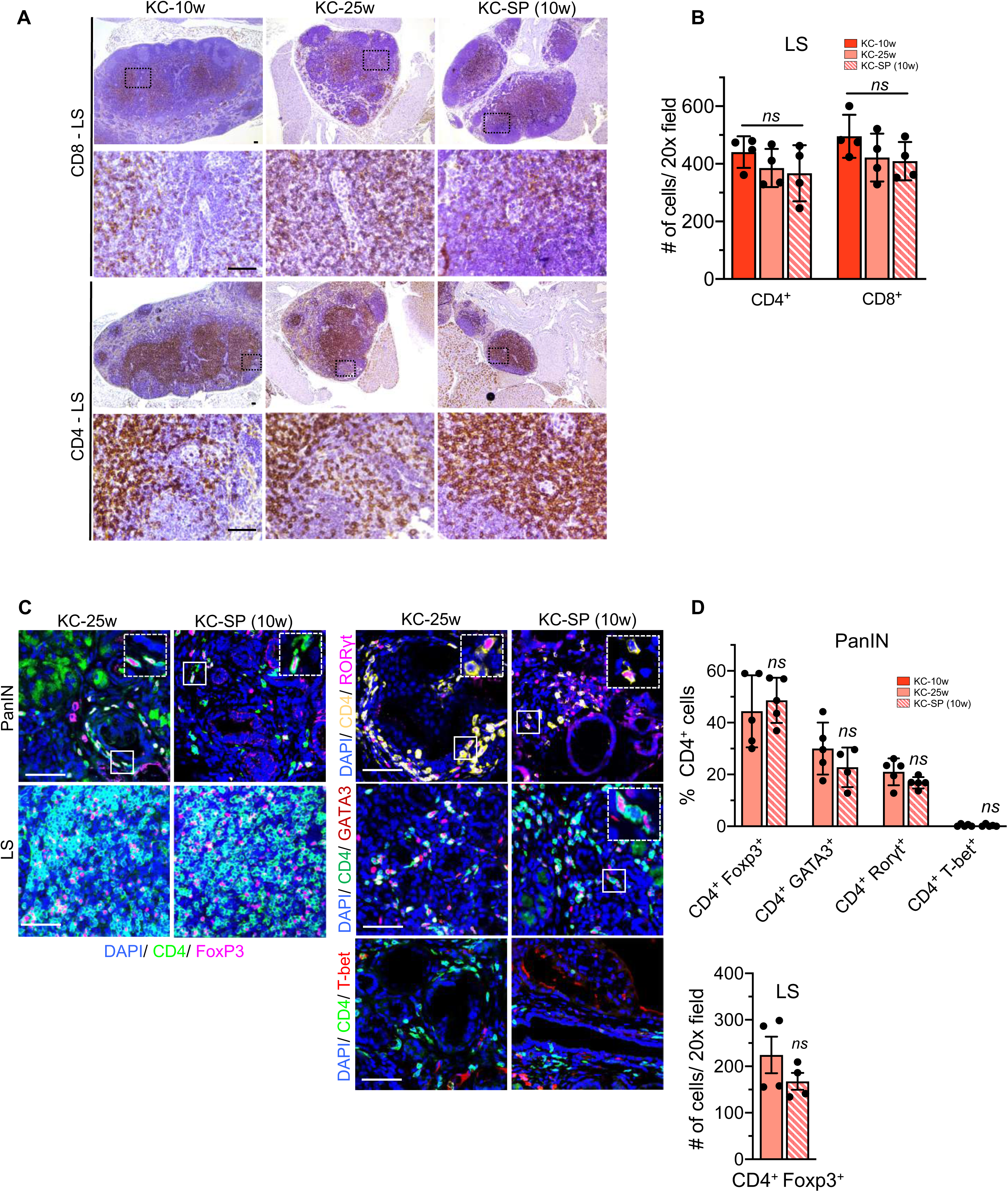
Analysis of T cell infiltrates in the lymphoid structures and PanIN lesions in KC mice with short-term pancreatitis. **Related to Figure 2.** (**A-B**) Representative CD4 and CD8 immunostaining (**A**), with quantification (**B**), of LS in the pancreata of KC-10w, KC-25w and KC-SP (10w) mice (n=4/ group). (**C-D**) Representative CD4^+^, Foxp3^+^ (**C**, left panel); CD4^+^, GATA3^+^; CD4^+^, RORγt^+^ and CD4^+^, γ T-bet^+^ (**C**, right panel), co-staining with quantification (**D**), in stage matched KC-25w and KC-SP (10w) mice pancreata (n=4-5/ group). In **B** and **D**, data are presented as mean ± SD. Significance was determined by one-way ANOVA with Dunnett’s multiple comparisons test in **B** and unpaired t-test in **D**. *ns*: not significant. Scale bars indicate 100 μm.

**Supplementary Figure 5:**
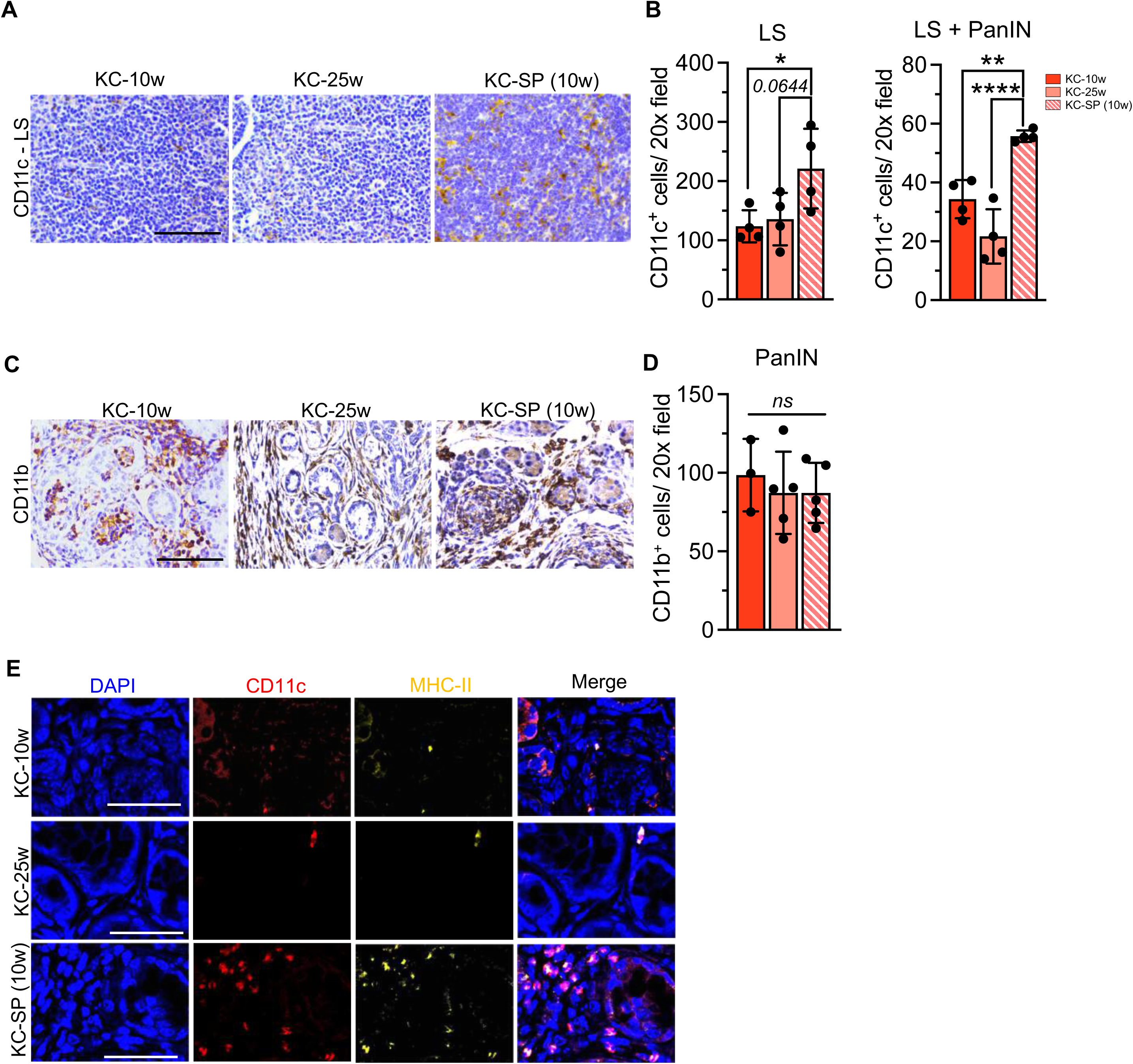
Analysis of myeloid and dendritic cell infiltration in lymphoid structures and PanIN lesions of KC mice with short-term pancreatitis. **Related to Figure 2.** (**A-B**) Representative CD11c immunostaining (**A**), with quantification (**B**), of LS in pancreata of KC-10w, KC-25w and KC-SP (10w) mice (n=4/ group). Note that the average number CD11c^+^ cells per 20x field in LS and PanIN lesions revealed higher CD11c^+^ cells in the KC-SP (10w) mice. (**C-D**) Representative CD11b immunostaining (**C**), with quantification (**D**), of PanIN lesions in KC-10w, KC-25w and KC-SP (10w) mice (n=3-5/ group). (**E**) Individual channel images from Figure 2B. In **B** and **D**, data are presented as mean ± SD. Significance was determined by one-way ANOVA with Dunnett’s multiple comparisons test in **B** and **D**. * P <0.05, ** P< 0.01, **** P <0.0001, *ns*: not significant. Scale bars indicate 100 μm.

**Supplementary Figure 6:**
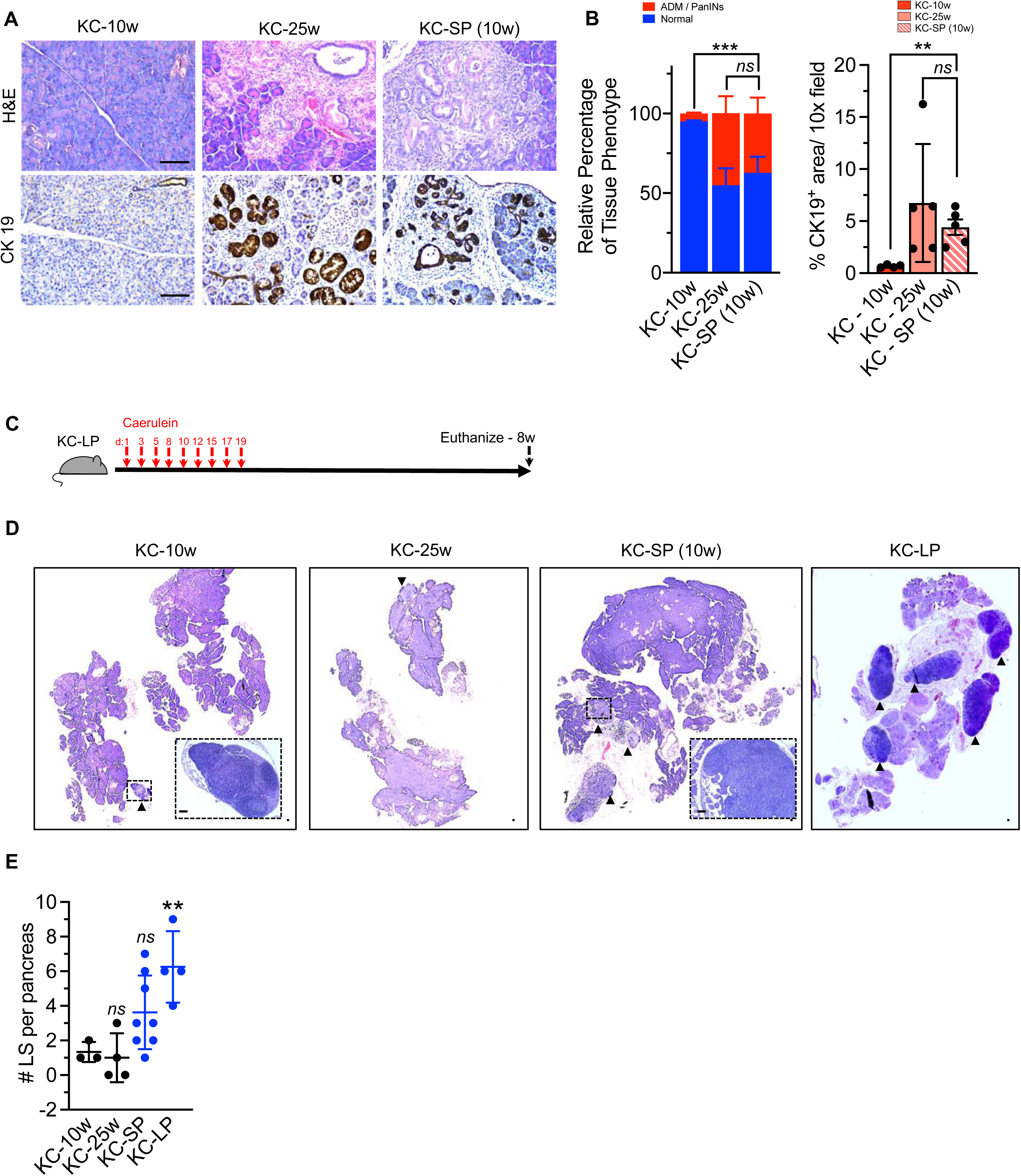
Pancreatitis accelerates tumor initiation and increases lymphoid structures in KC mice. **Related to Figure 2.** (**A-B**) Representative H&E and CK19 immunostaining (**A**) with quantification of PanIN lesions by H&E (**B**, left panel) and CK19 staining (**B**, right panel) of KC-10w (n=4), KC-25w (n=5) and KC-SP (10w) (n=5) mice pancreata. (**C**) Schematic representation of LP induction (each red arrow indicates injection of caerulein 4 times per day, 6 hours apart) in KC mice. (**D-E**) Representative H&E (**D**) with quantification of LS (**E**) in the pancreas of KC-10w (n=3), KC-25w (n=4), KC-SP (10w) (n=8) and KC-LP (n=4) mice. Note: Isotype treated KC-SP mice are included for this analysis. In **B, E** data are presented as mean ± SD. Significance was determined by two-way ANOVA with Tukey’s test for comparison of relative percentage of tissue phenotype and one-way ANOVA with Dunnett’s multiple comparisons test in **B** for immunostaining analysis and by one-way ANOVA with Dunnett’s multiple comparisons test in **E**. ** P <0.01, *** P <0.001, *ns*: not significant. Scale bars indicate 100 μm. Note: The KC-25w and KP-SP (10w) mice tissue in Supplementary Fig.S6A and S6B are repeated from Fig.3B, C and Fig. 3D, E to allow for direct comparisons.

**Supplementary Figure 7:**
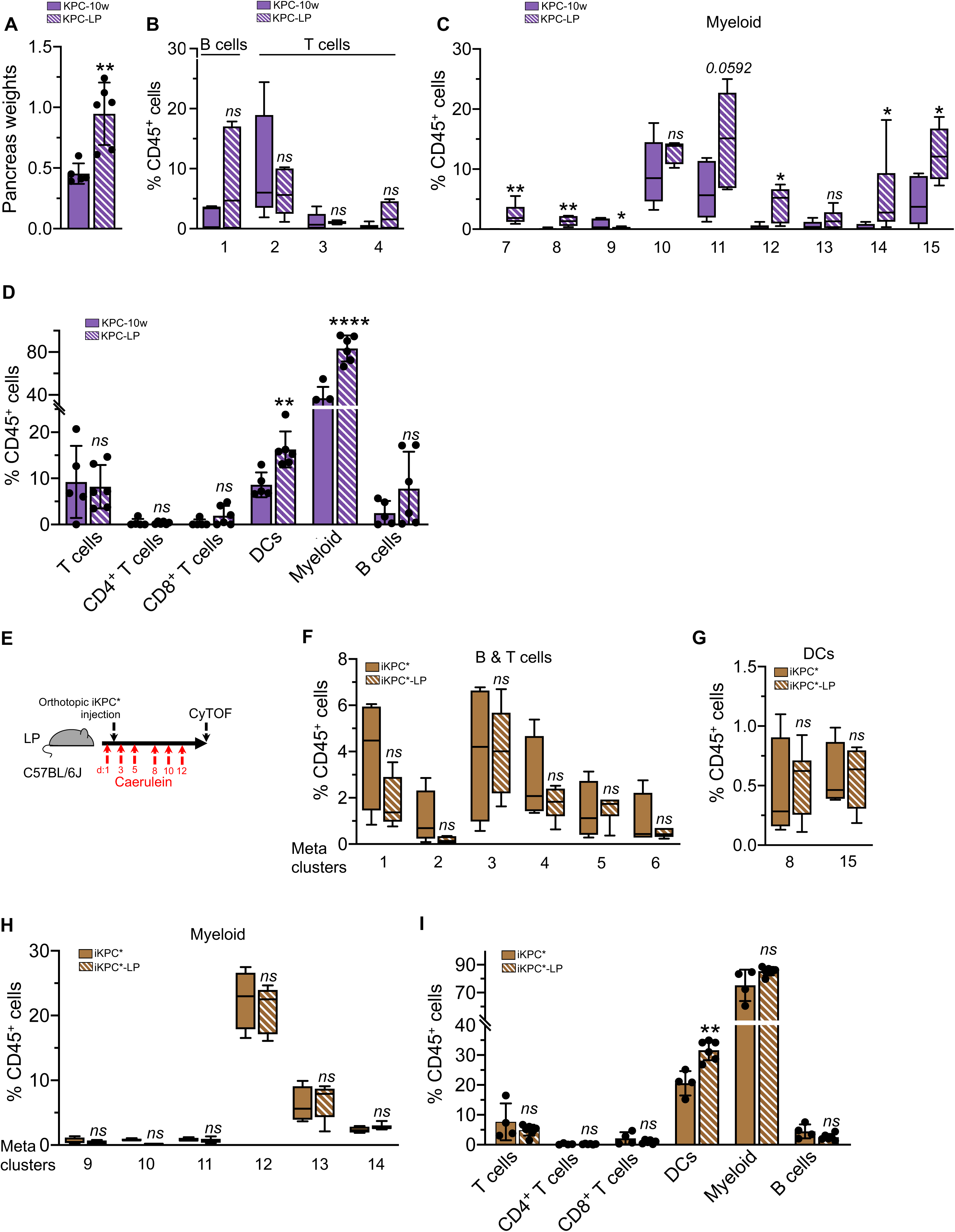
Additional CyTOF analysis of KPC and orthotopic iKPC* mice with long-term pancreatitis. **Related to Figure 2.** (**A**) Pancreas weights of KPC-10w (n=5) and KPC-LP (n=6) tumors. (**B-C**) Relative frequencies of B and T-cell MCs (**B**), and myeloid MCs (**C**) by unsupervised clustering of CyTOF markers in KPC-10w (n=5) and KPC-LP (n=6) mice pancreata. (**D**) Immune cell populations identified by Boolean gating of CyTOF markers in KPC-10w (n=5) and KPC-LP (n=6) tumors. (**E**) Schematic representation of LP-induction in orthotopic iKPC* mice. (**F-H**) Relative frequencies of MCs for indicated cell types. B and T cell MCs (**F**), DC MCs (**G**), and myeloid MCs (**H**), in iKPC* (n=4) and iKPC*-LP (n=6) tumors. (**I**) Immune cell populations identified by Boolean gating of CyTOF markers in iKPC* (n=4) and iKPC*-LP tumors (n=6). In **B, C, F, G** and **H** data are presented as box and whisker plots (min to max) and as mean ± SD in **A, D** and **I**. Significance was determined by unpaired T-test in **A**, Mann-Whitney test for comparison of MCs 1, 4 and unpaired T-test for comparison of MCs 2, 3 in **B**, Mann-Whitney test for comparison of MCs 7, 8, 10, 12, 14 and unpaired T-test for comparison of MCs 9, 11, 13, 15 in **C**, unpaired T-test for comparison of T cells, CD8^+^ T cells, myeloid, B cells and Mann-Whitney test for comparison of CD4^+^ T cells, DCs in **D**, unpaired T-test for comparison of MCs 1, 2, 3, 4 and Mann-Whitney test for comparison of MCs 5, 6 in **F**, unpaired T-test in **G and I**, Mann-Whitney test for comparison of MC 14 and unpaired T-test for all other comparisons in **H.** * P <0.05, ** P< 0.01, **** P <0.0001, *ns*: not significant.

**Supplementary Figure 8:**
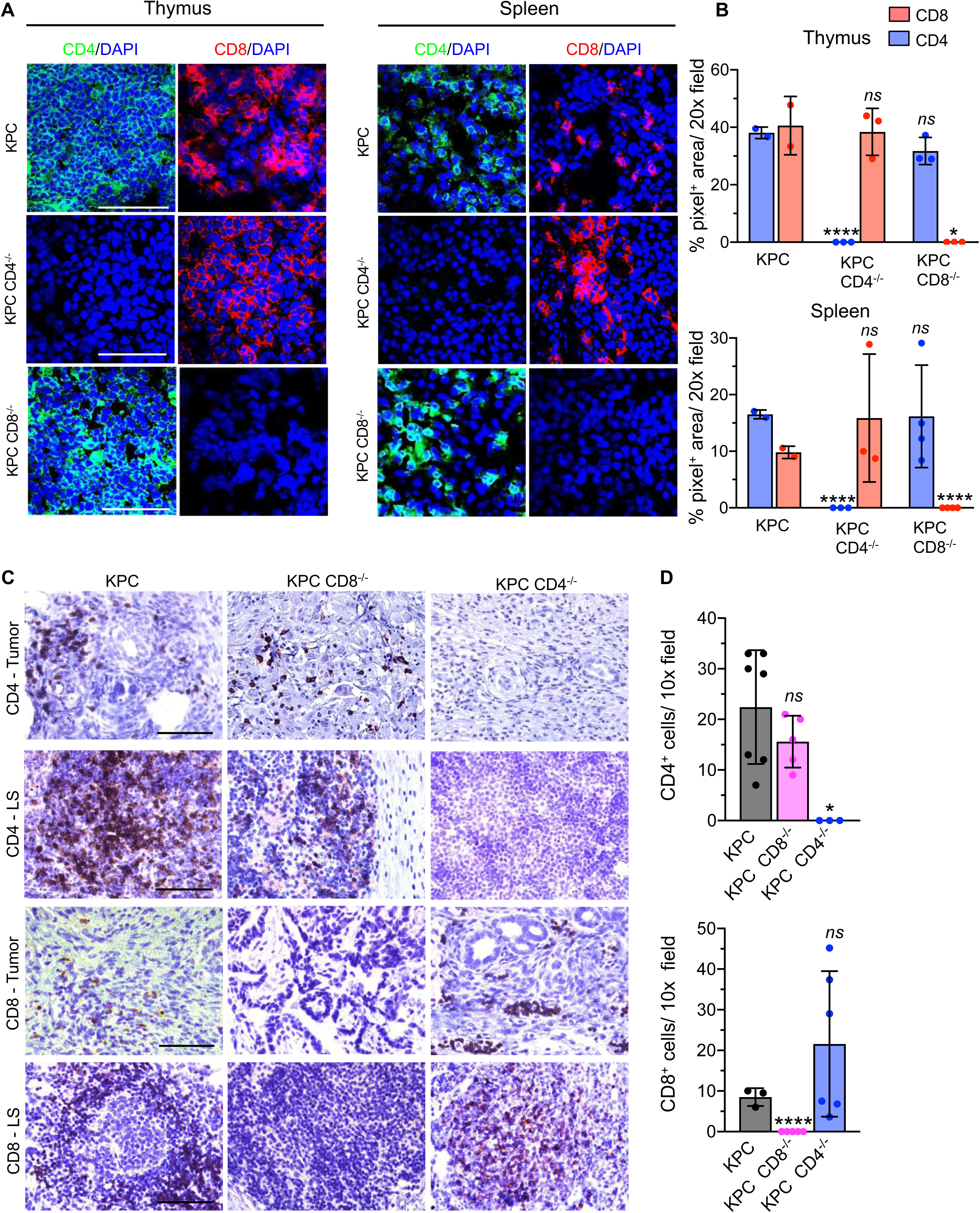
Immunolabeling of tumors and lymphoid organs of KPC, KPC CD4^-/-^ and KPC CD8^-/-^ mice for T cells. **Related to Figure 3.** (**A-B**) CD4 and CD8 immunostaining (**A**), with quantification (**B**), of percentage CD4^+^ or CD8^+^ area of thymus and spleen of KPC, KPC CD4^-/-^ and KPC CD8^-/-^ mice (n=2-4/ group). (**C-D**) CD4 and CD8 immunostaining (**C**), with quantification (**D**), of KPC, KPC CD4^-/-^ and KPC CD8^-/-^ tumors (n=3-7/ group). In **B** and **D**, data are presented as the mean ± SD. Significance was determined by unpaired t-test or Mann-Whitney test in **B and D**. *P<0.05, **** P<0.0001, *ns*: not significant. Scale bars indicate 100 μm. Note: Some data from this panel have been included in (Mahadevan et.al. 2023 under review) and are repeated here to provide confirmation of CD4^+^ and CD8^+^ T cells in the mouse models.

**Supplementary Figure 9:**
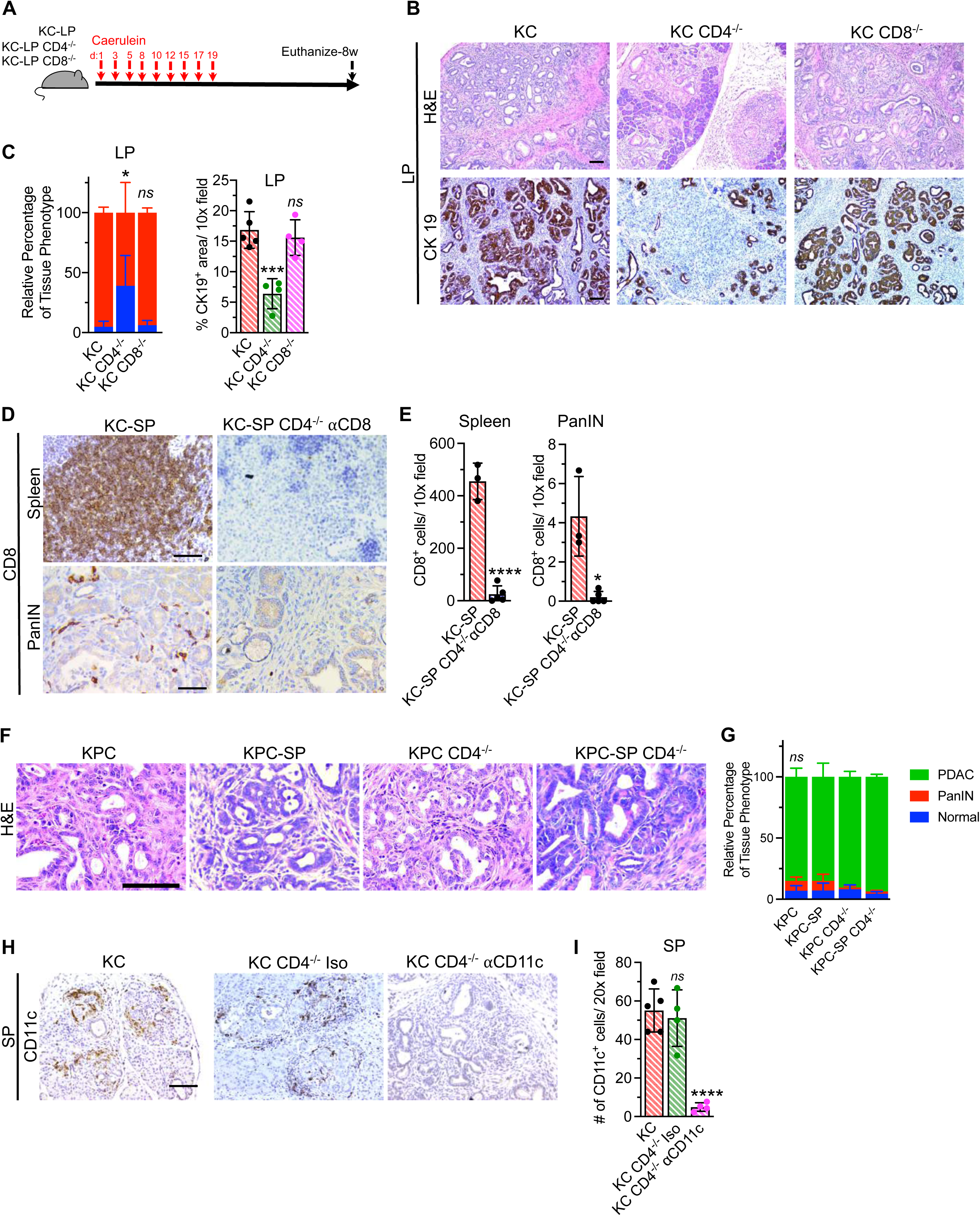
CD4^+^ T cells promote tumorigenesis and PDAC progression in KC and KPC mice with pancreatitis. **Related to Figure 3.** (**A**) Schematic representation of LP induction (each red arrow indicates injection of caerulein 4 times per day, 6 hours apart) in KC, KC CD4^-/-^ and KC CD8^-/-^ mice. (**B-C**) Representative H&E and CK19 immunostaining images (**B**), with quantification of PanIN lesions based on H&E (**C**, left panel) and CK19 staining (**C**, right panel) of KC-LP (n=5), KC-LP CD4^-/-^ (n=4), and KC-LP CD8^-/-^ (n=4) mice pancreata. (**D-E**) Representative CD8 immunostaining images (**D**), with quantification of CD8^+^ T cells in the spleen (**E**, left panel) and PanIN lesions (**E**, right panel) of KC-SP (n=3), KC-SP CD4^-/-^ ⍰CD8 (n=5) mice pancreata. Note: Same KC-SP control mice from Figure 2B. (**F-G**) Representative H&E images (**F**), with histological quantification (**G**) of endpoint KPC (n=5), KPC CD4^-/-^ (n=5) and KPC-SP (n=5) and KPC-SP CD4^-/-^ (n=7). (**H-I**) Representative CD11c immunostaining images (**H**), with quantification of CD11c^+^ cells in PanINs of KC-SP (n=5), KC-SP CD4^-/-^ Iso (n=4), KC-SP CD4^-/-^ ⍰CD11c (n=4) mice pancreata (**I**). Note: Same KC-SP control mice from Figure 2B. In **C, E, G** and **I** data are presented as mean ± SD. Significance was determined by two-way ANOVA with Tukey’s test for comparison of relative percentage of tissue phenotype in **C** and **G**, one-way ANOVA with Dunnett’s multiple comparisons test for immunostaining analysis in **C** and **I**, unpaired T-test for comparison of CD8 quantification in the spleen in **E**, Mann-Whitney test for comparison of CD8 quantification in PanINs in **E**. *P<0.05, *** P < 0.001, **** P <0.0001, *ns*: not significant. Scale bars indicate 100μm.

**Supplementary Figure 10:**
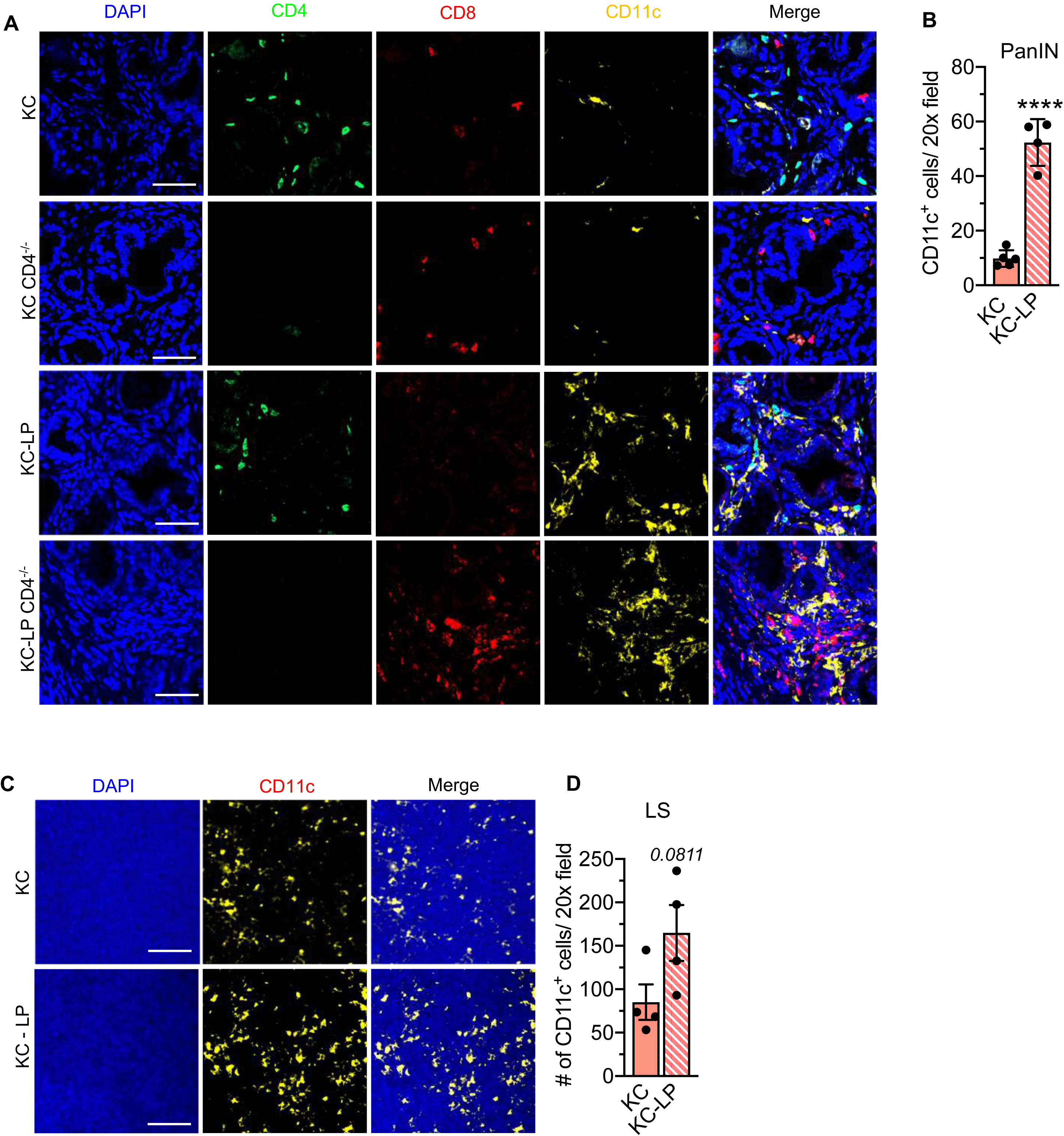
Individual channel images from main figure and additional analysis of CD11c staining in KC mice. **Related to Figure 3.** (**A-B**) Individual channel images (**A**), with quantification of CD11c^+^ cells in KC (n=5) and KC-LP (n=4) mice (**B**) from Figure 3I. (**C-D**) Representative CD11c immunostaining (**C**), with quantification of CD11c^+^ cells in the LS (**D**) in the pancreas of KC and KC-LP mice (n=4/ group). Significance was determined by unpaired t-test in **B** and **D**. **** P<0.0001. Scale bars indicate 100 μm.

**Supplementary Figure 11:**
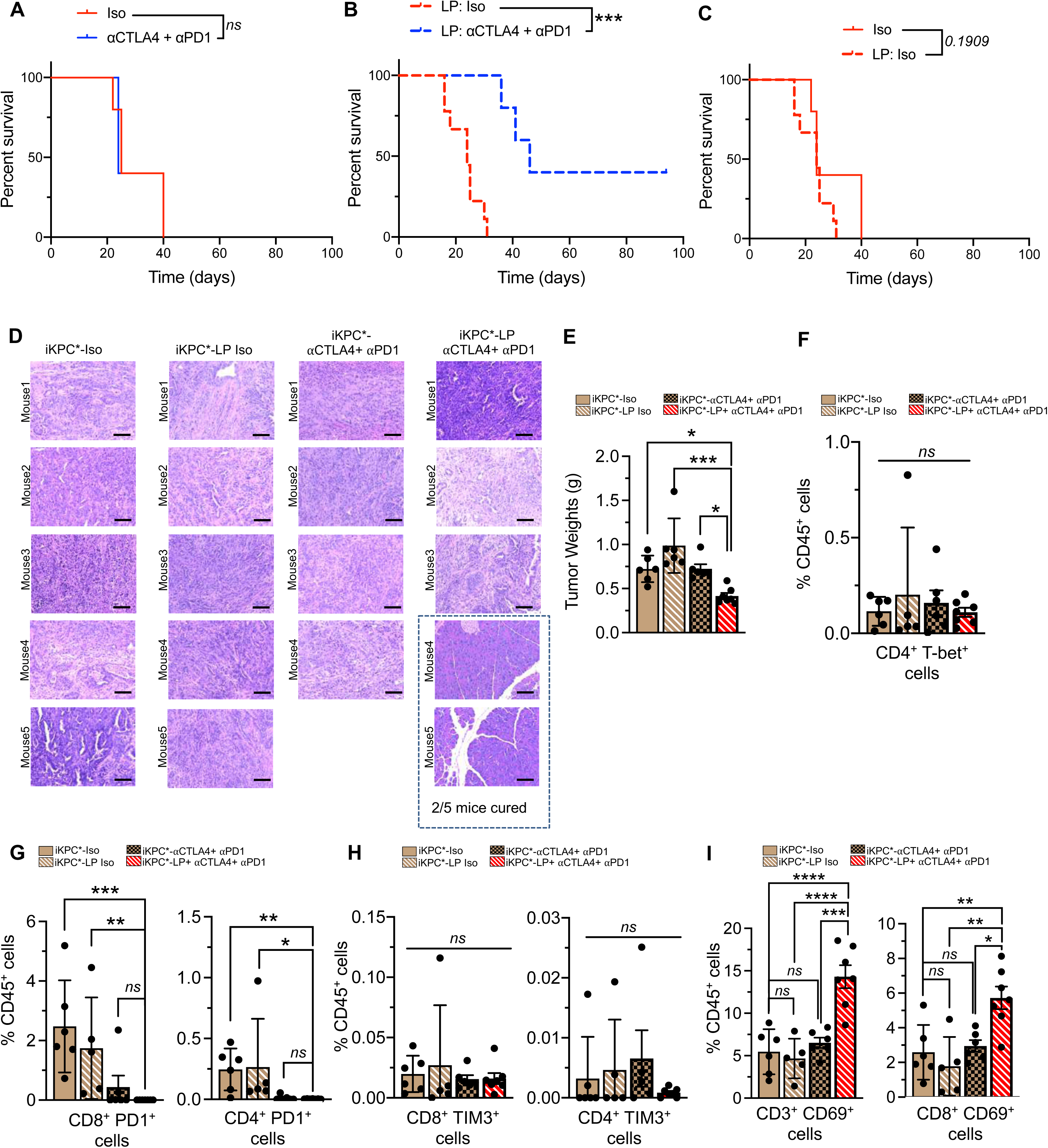
Individual survival curves and additional analysis of tumor infiltrating lymphocytes from main figure. **Related to Figure 4.** (**A-C**) Individual Kaplan Meier survival curves of indicated groups of orthotopic iKPC* tumor mice from Figure 4E. (**D**) Representative images of iKPC* mice orthotopic tumors or pancreas of indicated experimental groups at autopsy. (n=4-5/ group). (**E**) Tumor weights of isotype (n=6), LP-Isotype (n=6), ⍰CTLA4 + ⍰PD1 (n=6) and LP-⍰CTLA4 + ⍰PD1 (n=7) treated mice euthanized on day 14 following orthotopic iKPC* injection. (**F-H**) Immunophenotyping analysis of exhaustion or activation markers on T cells of indicated groups. CD4^+^ T-bet^+^ (Th_1_) cells (**F**), CD8^+^PD1^+^ and CD4^+^PD1^+^ cells (**G**), and CD4^+^TIM3^+^ and CD8^+^TIM3^+^ cells (**H**) measured as a percentage of CD45^+^ cells. (**I**) Immunophenotyping analysis of activation and memory marker CD69 on CD3^+^ and CD8^+^ T cells as a percentage of CD45^+^ cells. iKPC*-Iso (n=6), iKPC*-LP Iso (n=5), iKPC*-⍰CTLA4 + ⍰PD1 (n=6) and iKPC*-LP + ⍰CTLA4 + ⍰PD1 (n=7) in **F-I**. In (**E-I**), data are presented as mean ± SD. Significance was determined by Kruskal-Wallis with Dunn’s multiple comparisons test in **E, F, G** and **H**, one-way ANOVA with Sidak’s multiple comparisons test in **I**. *P <0.05, **P <0.01, *** P <0.001, **** P<0.0001, *ns*: not significant.

**Supplementary Figure 12:**
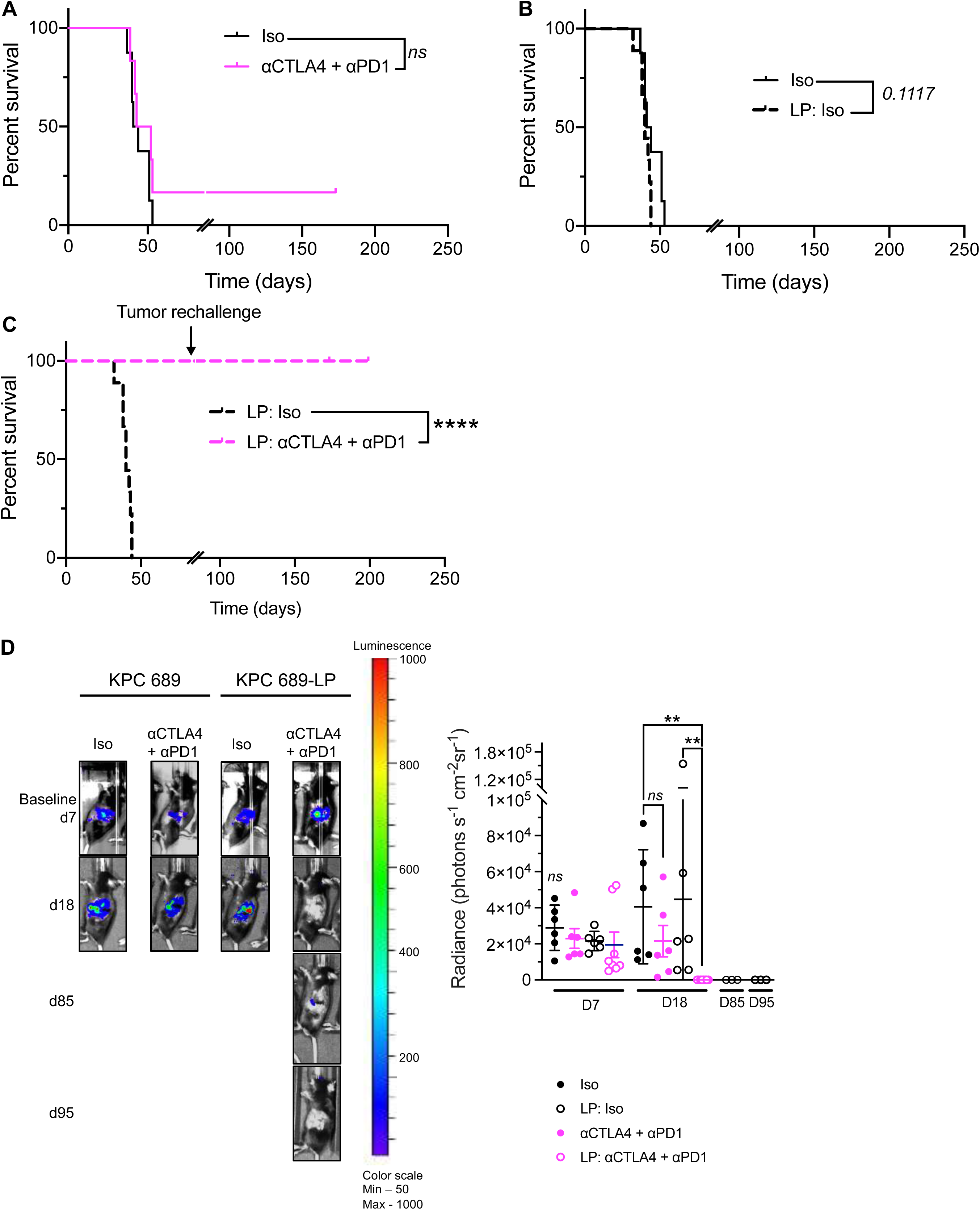
Individual survival curves and IVIS images from main figure. **Related to Figure 4.** (**A-C**) Individual Kaplan Meier survival curves of indicated groups of orthotopic KPC689 tumor bearing mice from Figure 4I. (**D**) Representative IVIS imaging (left panel) and quantification (right panel) of baseline on day 7, follow up on day 18, imaging on day 85 before re-challenge and on day 95 in indicated experimental groups. Significance was determined by log-rank test in **A-C**, Kruskal-Wallis with Dunn’s multiple comparisons test in **D**.**P<0.01, ****P <0.0001, *ns*: not significant.

**Supplementary Figure 13:**
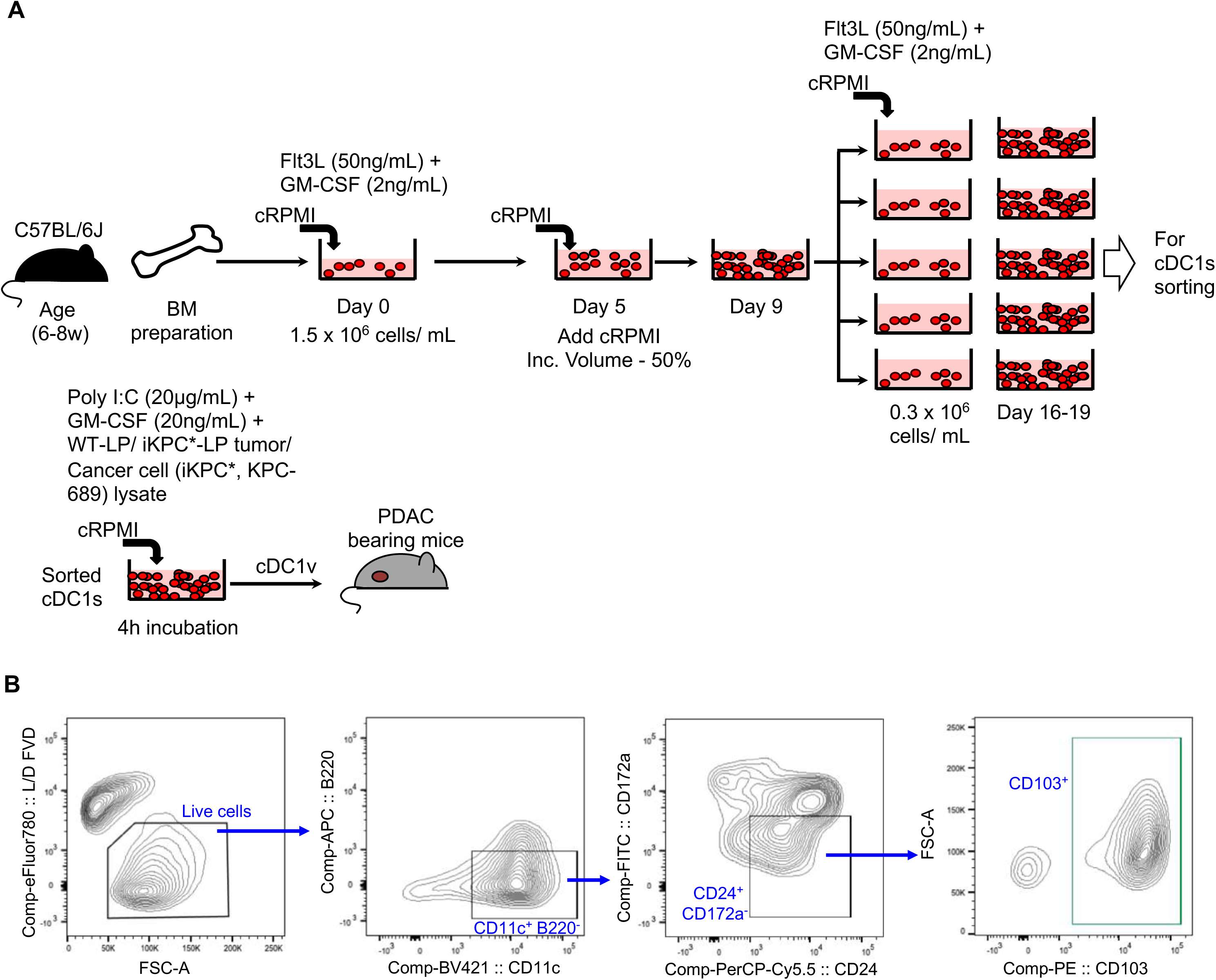
Schematic for cDC1 vaccine preparation from bone marrow culture and gating strategy. **Related to Figure 5.** (**A**) Schematic for generation of cDC1 vaccine from bone marrow culture of 6-8w old C57BL/6J mice. (**B**) Gating strategy for sorting cDC1 population (Live/dead^-^, CD11c^+^B220^-^CD24^+^CD172a^-^ CD103^+^ cDC1s).

**Supplementary Figure 14:**
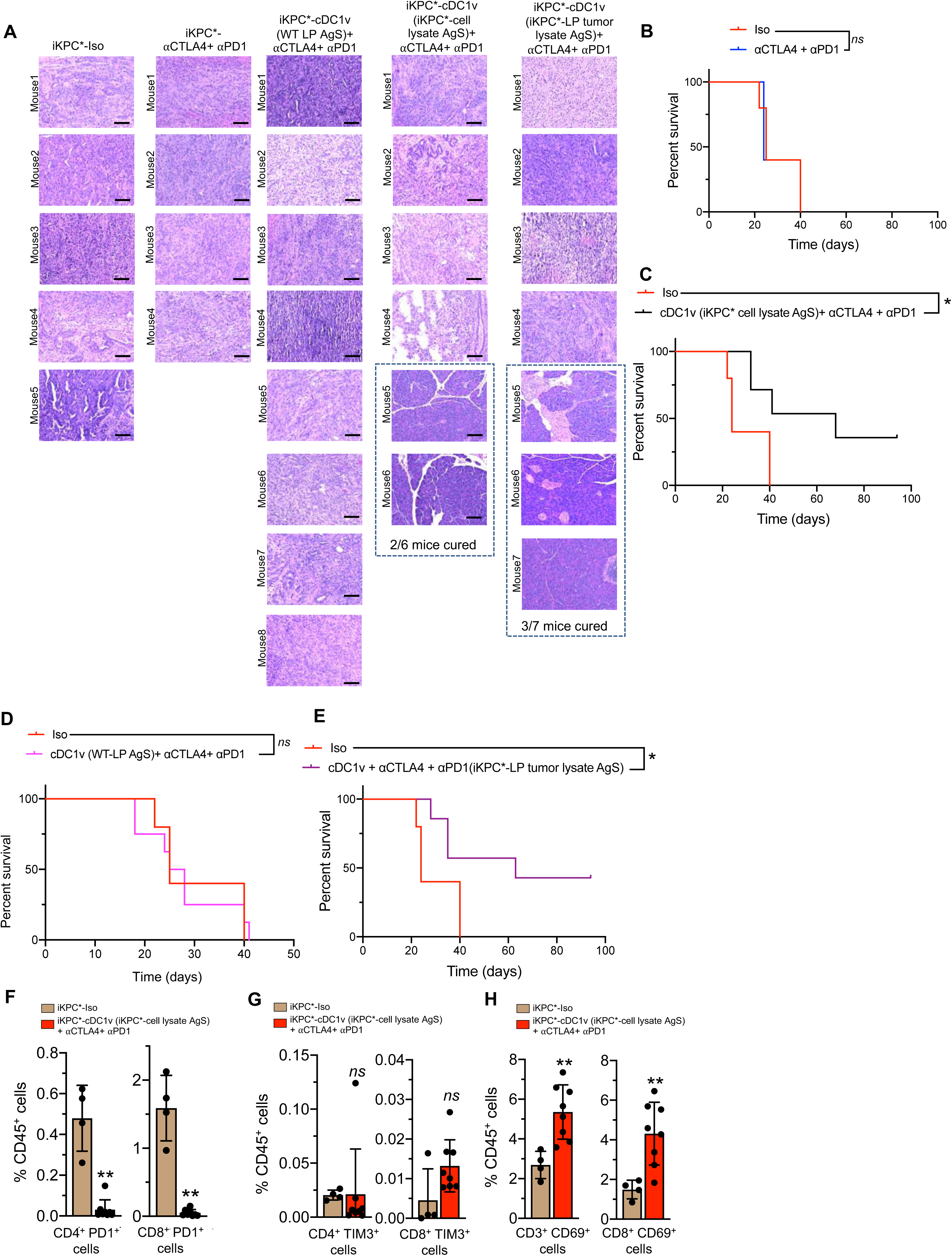
cDC1 vaccine sensitizes the orthotopic iKPC* tumors to checkpoint blockade immunotherapy. **Related to Figure 5.** (**A**) Representative images of iKPC* mice orthotopic tumors or pancreas of indicated experimental groups at autopsy. (n=4-8/ group). Note: KPC-Iso H&E is repeated from S11D. (**B-E**) Individual Kaplan Meier survival curves of indicated groups of orthotopic iKPC* tumor mice from Figure 5B. (**F-G**) Immunophenotyping analysis of exhaustion markers on T cells. CD4^+^PD1^+^ and CD8^+^PD1^+^ cells (**F**); and CD4^+^TIM3^+^ and CD8^+^TIM3^+^ cells (**G**) measured as a percentage of CD45^+^ cells. (**H**) Immunophenotyping analysis of activation and memory marker CD69 on CD3^+^ and CD8^+^ T cells as a percentage of CD45^+^ cells. In (**F-H**), data are presented as mean ± SD. Significance was determined by unpaired t-test in **H**, Mann-Whitney test in **F and G**, and by log-rank test in **B-E**. *P <0.05, **P <0.01, *ns*: not significant. Scale bars indicate 100 μm. Note: Supplementary Fig. 14B is repeated from Supplementary Fig. 11A

**Supplementary Figure 15:**
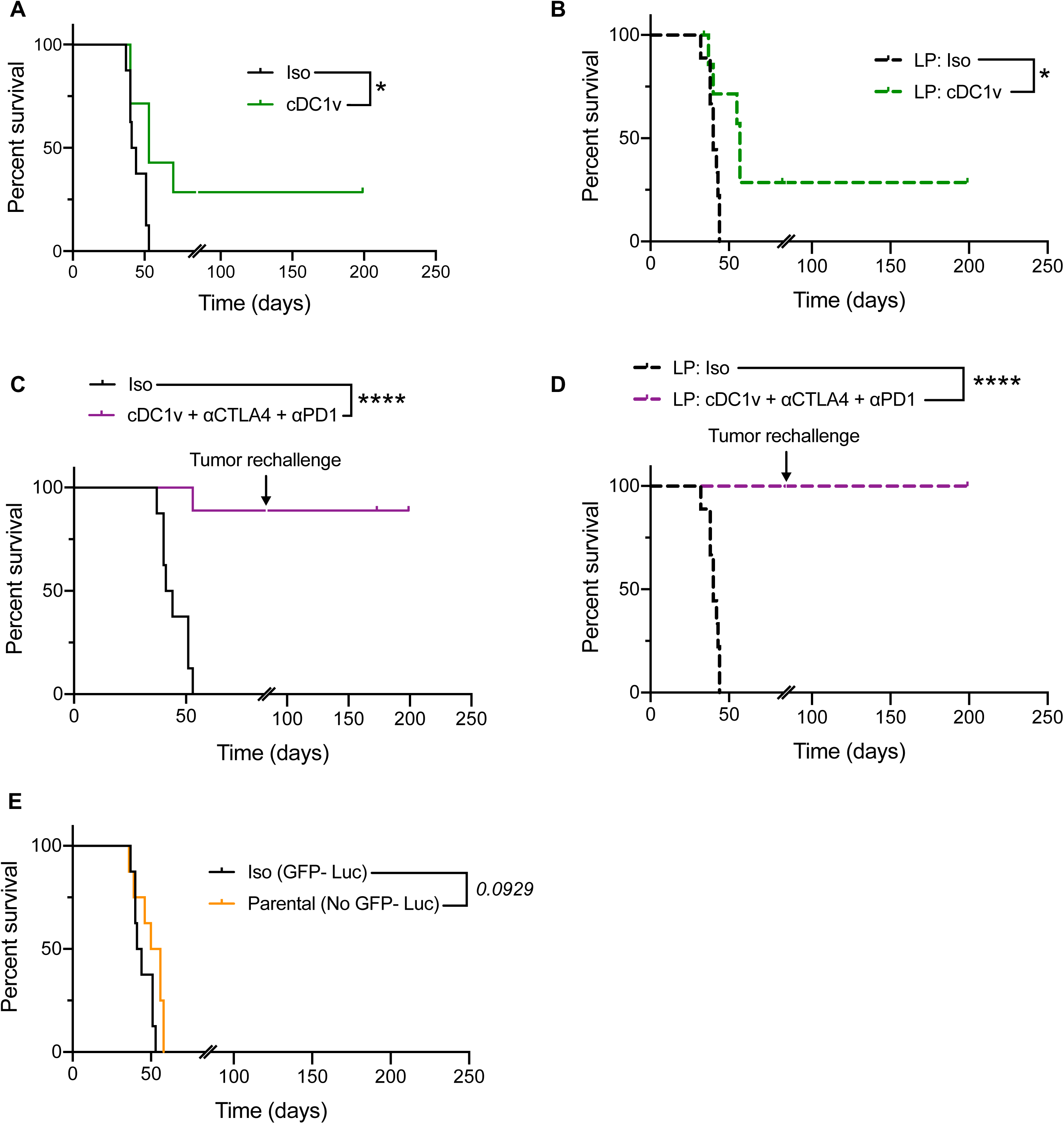
(**A-E**) **Individual survival curves from main figure. Related to** Figure 5. (**A-E**) Individual Kaplan Meier survival curves of indicated groups of orthotopic KPC689 tumor bearing mice from Figure 5H. *P <0.05, ****P <0.0001, *ns*: not significant. Note: S15A is repeated from S12A.

**Supplementary Figure 16:**
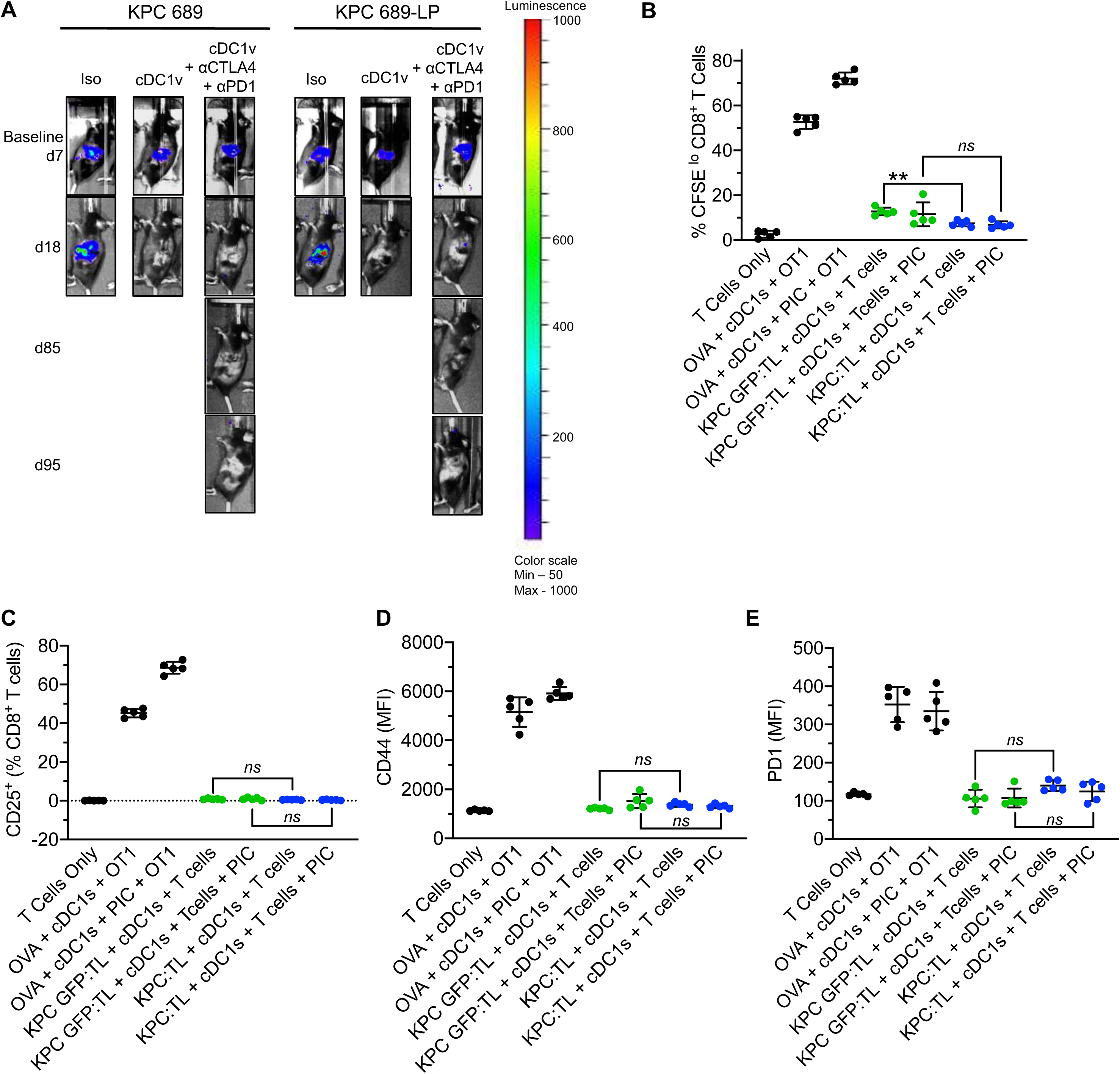
Additional analysis on the effect of GFP-Luc on T cell response and cDC1 vaccine treated KPC-689 orthotopic tumor bearing mice. **Related to Figure 5.** (**A**) Representative IVIS imaging of baseline on day 7, follow up on day 18, imaging on day 85 before re-challenge and on day 95 in indicated experimental groups. (**B-E**) KPC-689 GFP-Luc containing and parental tumor cell lysates demonstrated similar levels of T cell activation. Bone marrow-derived CD103^+^ cDC1s were stimulated with poly I:C and/or tumor cell lysates of KPC GFP-expressing cells or parental KPC tumors. Cells were co-cultured with CFSE-labelled wild type CD8^+^ T cells with cDC1s. OT-I splenic CD8^+^ T cells were cocultured with CD103^+^ cDC1s stimulated with ovalbumin and/or poly I:C as a positive control. % CFSE^lo^, CD8^+^ T cells (**B**), CD25^+^ (% CD8^+^ T cells) (**C**), CD44 (MFI) (**D**), and PD1 (MFI) (**E**), on T cells was quantified via flow cytometry (n=5/ group). In **B-E**, data are presented as mean ± SD., unpaired T-test in **B, C** and **D**, and Mann-Whitney test in **E**. **P <0.01, ***P <0.001, *ns*: not significant. Note: Iso groups in Supplementary Fig. 16A is repeated from Supplementary Fig. 12D. to allow for direct comparisons

**Supplementary Figure 17:**
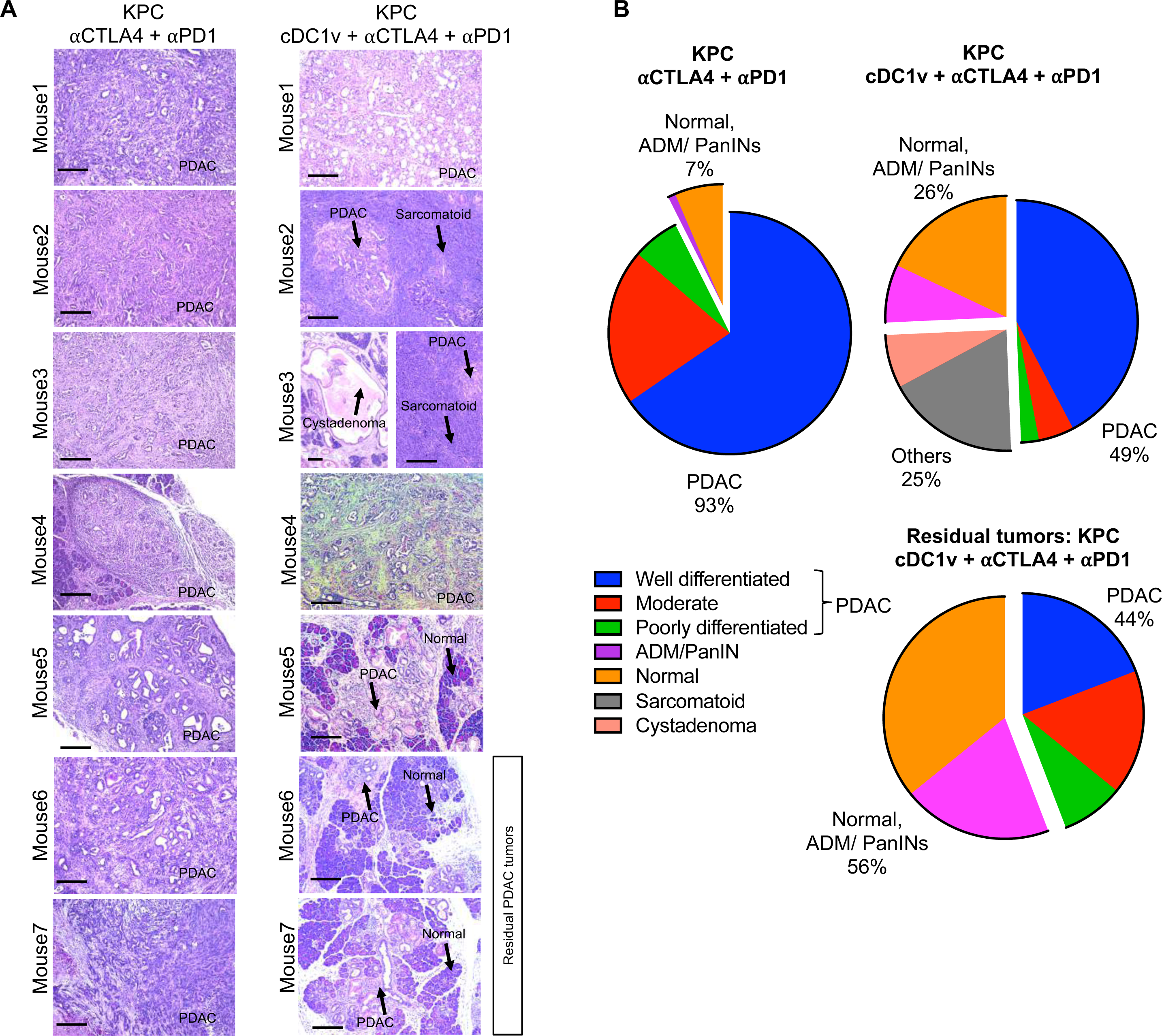
Additional analysis of KPC tumors treated with cDC1 vaccine and checkpoint blockade therapy. **Related to Figure 6.** (**A-B**) Representative H&E images (**A**), with pie-chart showing quantification of tissue histology in ⍰CTLA4 + ⍰PD1 (n=7) and cDC1v + ⍰CTLA4 + ⍰PD1 (n=7) KPC mice from Figure 5K (**B**). Data are presented as pie chart in **B**. Scale bars indicate 100 μm.

**Supplementary Figure 18:**
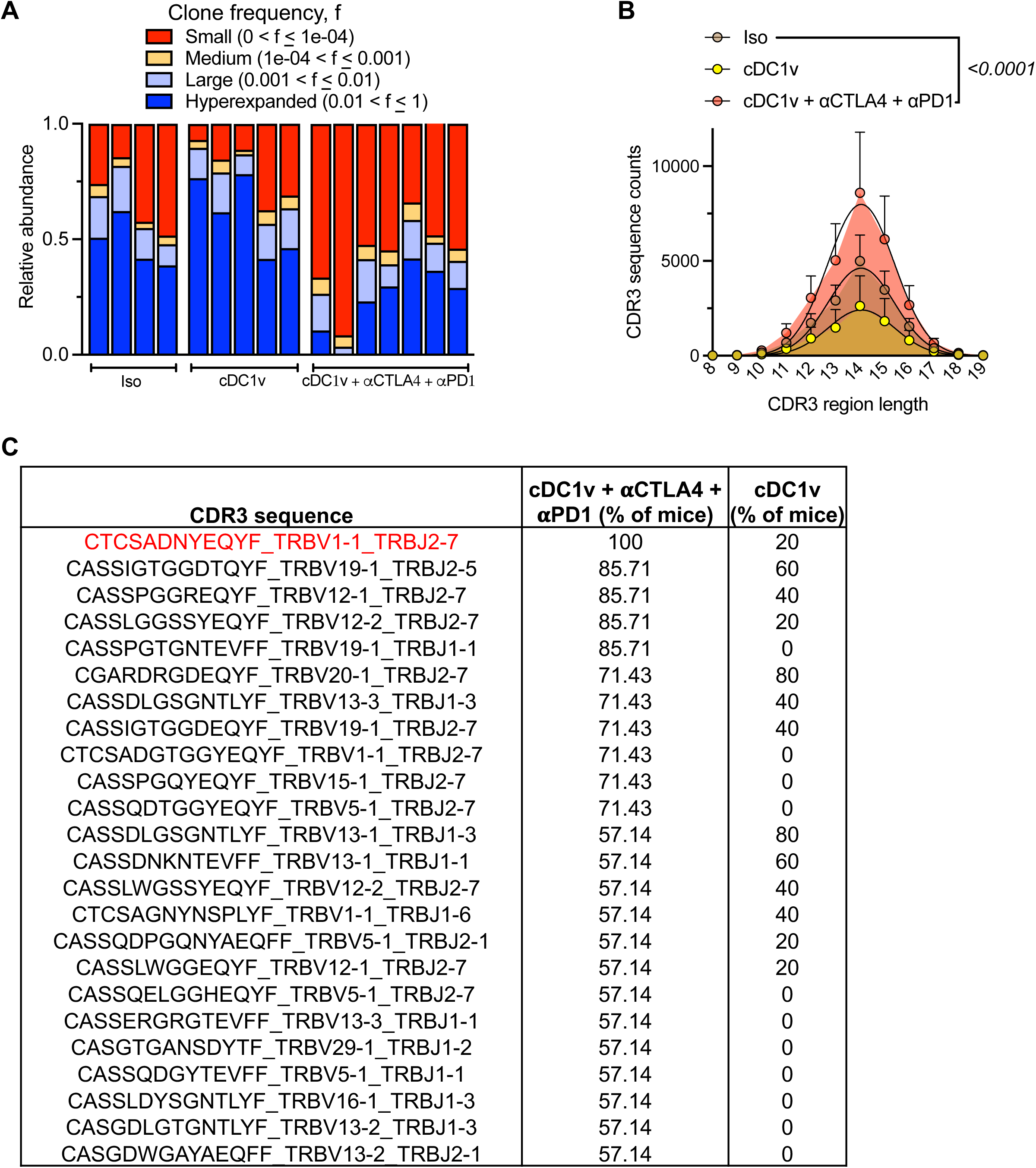
Additional TCR sequencing analysis of tumor infiltrating CD8^+^ T cells in orthotopic iKPC* model. **Related to Figure 7.** (**A**) Relative abundance of small, medium, large and hyperexpanded clones based on clone frequency. (**B**) CDR3 region lengths of tumor infiltrating CD8^+^ T cells in Iso (n=4), cDC1v (n=5) and cDC1v + ⍰CTLA4 + ⍰PD1 (n=7) treated mice. (**C**) List of unique CDR3 AA sequences enriched in >50% of cDC1v + ⍰CTLA4 + ⍰PD1 treated mice in descending order of frequencies. In **A**, data are presented as mean and as mean ± SD in **B**.

**Supplementary Figure 19:**
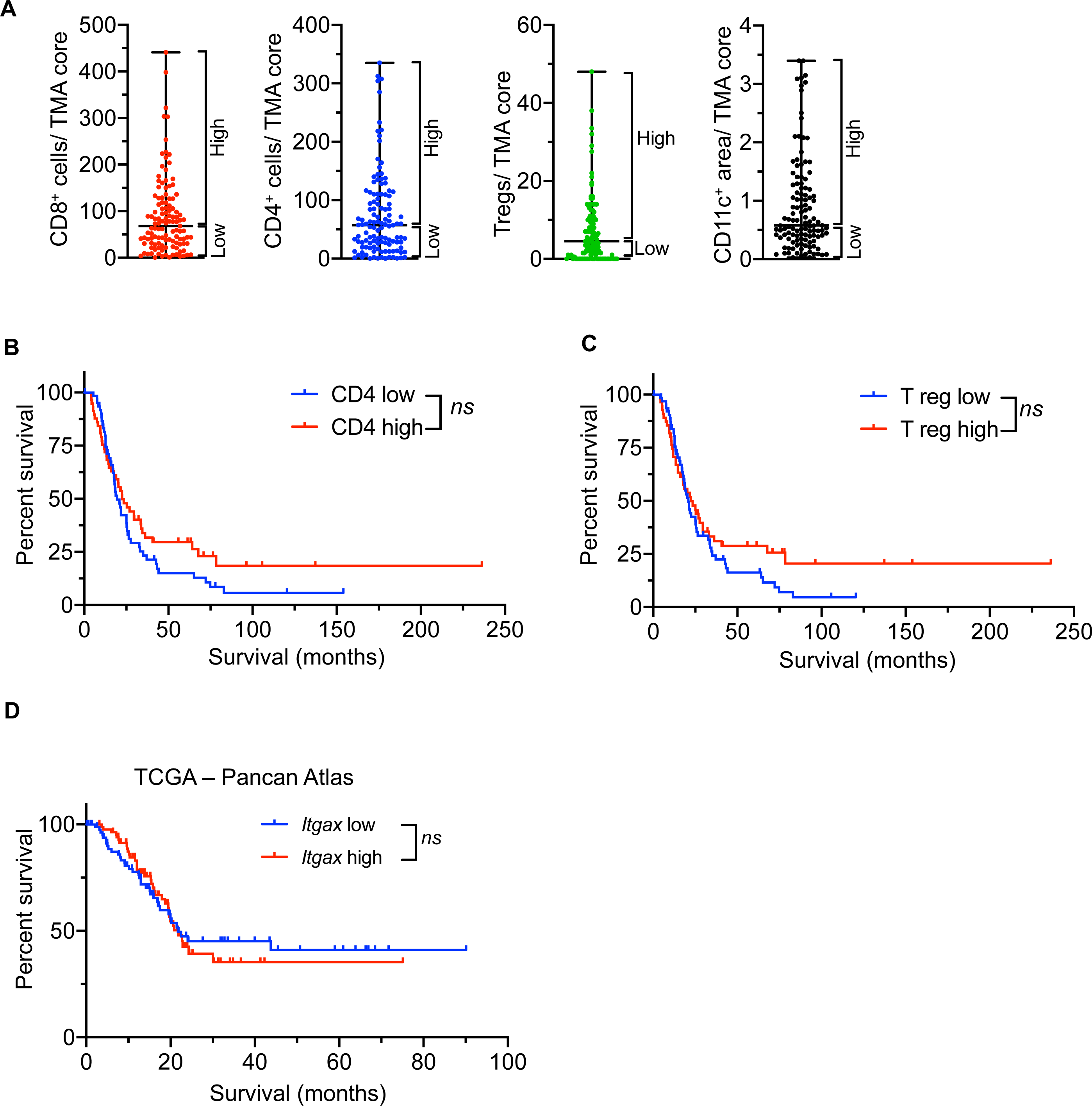
Additional analysis of the role of immune cell infiltration in human PDAC. **Related to Figure 8.** (**A**) Stratification of human PDAC as ‘high’ and ‘low’ for each cell type based on median value among all patient samples. (**B-C**) Kaplan-Meier survival curves for disease specific survival (DSS) of CD4 high (n=58) vs. CD4 low (n=62) tumors (**B**), and Treg high (n=55) Vs. Treg low (n=65) tumors (**C**). (**D**) Kaplan-Meier survival curves for DSS of PDAC patients with *Itgax* (CD11c) high (n=86) vs. low (n=87) expression based on RNA seq analysis in samples from TCGA-PanCanAtlas dataset. *ns*: not significant.

