## Supplementary Tables 1-5 for "Antigen-presenting type-I conventional dendritic cells facilitate curative checkpoint blockade immunotherapy in pancreatic cancer"

**Supplementary Table 1: Markers for cDC1 sorting and analysis of $CD8^+$ T cell stimulation experiments**

**cDC1 sorting**

| Marker | Fluorophore | Dilution | Cat # (Clone) |
|---|---|---|---|
| CD11c | BV421 | 1:200 | Tonbo Biosciences 20-0114 (N418) |
| CD45R | APC | 1:200 | Biolegend 103212 (RA3-6B2) |
| CD24 | PerCPCy5.5 | 1:200 | Biolegend 101824 (M1/69) |
| CD172a | FITC | 1:200 | Biolegend 144006 (P84) |
| CD103 | PE | 1:200 | Biolegend 121406 (2E7) |
| CD3 | PE-Cy7 | 1:200 | Biolegend 300419 (17A2) |

**$CD8^+$ T cell stimulation experiments**

| Marker | Fluorophore | Dilution | Cat # (Clone) |
|---|---|---|---|
| CD44 | APC | 1:200 | Biolegend 103012 (IM7) |
| CD103 | BV605 | 1:200 | Biolegend 121433 (2E7) |
| CD11c | BV650 | 1:200 | BD Horizon 564079 (HL3) |
| CD3 | PerCPCy5.5 | 1:200 | Biolegend 100218 (17A2) |
| PD1 | PeCy7 | 1:200 | Biolegend 109110 (29F.IAI2) |
| CD8 | BV711 | 1:200 | Biolegend 344734 (53-6.7) |
| CD25 | PE | 1:200 | Tonbo Biosciences 50-0251 (PC61.5) |

### Supplementary Table 2: Antibodies used for TSA staining

| Antigen | Antigen retrieval buffer | Primary antibody | | | Secondary antibody | | TSA fluorophore |
|---|---|---|---|---|---|---|---|
| | | Vendor | Catalog # | Dilution | Vendor | Polymer | |
| **Mouse tissue staining** | | | | | | | |
| CD11c | Citrate (pH=6) | Cell Signaling Technology | 97585S | 1:350 | BioCare | Rabbit-on-Rodent HRP | Opal 650 |
| MHC-II | Citrate (pH=6) | Thermo Fisher Scientific | 14-5321-82 | 1:100 | BioCare | Rat HRP | Opal 520 |
| CD4 | Tris-EDTA (pH=9) | Abcam | ab183685 | 1:400 | BioCare | Rabbit-on-Rodent HRP | Opal 520 |
| CD8 | Tris-EDTA (pH=9) | Cell Signaling Technology | 98941s | 1:250 | BioCare | Rabbit-on-Rodent HRP | Opal 570 |
| Foxp3 | Tris-EDTA (pH=9) | ThermoFischer Scientific | 14-4771-80 | 1:50 | BioCare | Rat HRP | Opal 650 |
| GATA3 | Tris-EDTA (pH=9) | Cell Signaling Technology | 5852S | 1:1500 | BioCare | Rabbit-on-Rodent HRP | Opal 650 |
| Rorγt | Tris-EDTA (pH=9) | Abcam | ab207082 | 1:1000 | BioCare | Rabbit-on-Rodent HRP | Opal 650 |
| T-bet | Tris-EDTA (pH=9) | Abcam | ab91109 | 1:100 | Vector Laborotaries | M.O.M. Immunodetection kit | Opal 650 |
| **Human tissue PDAC-TMA staining** | | | | | | | |
| CD4 | Tris-EDTA (pH=9) | Biocare | CM153CK | 1: 50 | BioCare | MACH2 Mouse HRP | Opal 520 |
| CD8 | Tris-EDTA (pH=9) | Dako | M7103 | 1:100 | BioCare | MACH2 Mouse HRP | Opal 570 |
| Foxp3 | Tris-EDTA (pH=9) | Abcam | ab20034 | 1:50 | BioCare | MACH2 Mouse HRP | Opal 650 |

**Supplementary Table 3: T cell and DC panel flowcytometry antibodies for immunotyping**

**T cell panel**

| Marker | Fluorophore | Dilution | Cat # (Clone) |
|---|---|---|---|
| Granzyme B | APC | 1:100 | Invitrogen GRB05 (GB11) |
| CD8 | BV650 | 1:200 | Biolegend 100742 (53-6.7) |
| CD45 | Pacific Blue | 1:100 | Biolegend 103126 (30-F11) |
| Live/dead | eFluor780 (APC-Cy7) | 1:1000 | eBiosciences 65-0865-14 |
| CD11b | BV711 | 1:400 | BD 563168 (M1/70) |
| Ki67 | Alexa488 | 1:100 | BD 558616 (B56) |
| CD49b | PE | 1:100 | eBiosciences 12-5971-81 (Dx5) |
| CD4 | BV605 | 1:200 | Biolegend 100548 (RM4-5) |
| Foxp3 | Alexa700 | 1:50 | eBiosciences 56-5773-82 (FJK-16s) |
| CD3 | PE-Cy7 | 1:200 | eBiosciences 25-0031-82 (145-2c11) |

**T cell activation/exhaustion panel**

| Marker | Fluorophore | Dilution | Cat # (Clone) |
|---|---|---|---|
| CD8 | BV650 | 1:200 | Biolegend 100742 (53-6.7) |
| CD45 | Pacific Blue | 1:100 | Biolegend 103126 (30-F11) |
| Live/dead | eFluor 780 (APC-Cy7) | 1:1000 | eBiosciences 65-0865-14 |
| CD69 | PE | 1:100 | Invitrogen 12-0691-81 |
| CD4 | BV605 | 1:200 | Biolegend 100548 (RM4-5) |
| CD3 | Alexa700 | 1:50 | eBiosciences 56-0032-82 (17A2) |
| CD62L | APC | 1:100 | BD 561919 (MEL-14) |
| PD1 | PerCp-Cy5.5 | 1:100 | Biolegend 135208 (29F.1A12) |
| TIM3 | PE/Dizzle (PE-TxR) | 1:100 | Biolegend 134013 (B8.2C12) |
| T-bet | PE-Cy7 | 1:100 | eBioscience 25-5825-80 (eBio4B10) |

**DC panel**

| Marker | Fluorophore | Dilution | Cat # (Clone) |
|---|---|---|---|
| CD45 | PB | 1:100 | Biolegend 103126 (30-F11) |
| Ly-6G | PE-Cy7 | 1:200 | BD 560601 (1A8) |
| CD11b | BV711 | 1:400 | BD 563168 (M1/70) |
| CD11c | PE-CF594 | 1:100 | BD 562454 (HL3) |
| Live/dead | eFluor 780 (APC-Cy7) | 1:1000 | eBioscience 65-0865-14 |
| CD40 | BV650 | 1:100 | BD 740492 (3/23) |
| CD86 | BV605 | 1:100 | BD 563055 (GL1) |
| Ly-6C | APC** | 1:100 | BD 560595 (AL-21) |

| | | | |
|---|---|---|---|
| CD3 | A700 | 1:50 | eBioscience 56-0032-82 (17A2) |

**DC subset panel**

| Marker | Fluorophore | Dilution | Cat # (Clone) |
|---|---|---|---|
| B220 | BV711 | 1:100 | Biolegend 103211 (RA3-6b2) |
| CD11c | PE-CF594 | 1:100 | BD 562454 (HL3) |
| CD11b | BV786 | 1:200 | BD 740861 (M1/70) |
| CD172a | FITC | 1:100 | Biolegend 144005 (P84) |
| Live/dead | BV510 | 1:1000 | BD 564406 |
| XCR1 | BV421 | 1:100 | Biolegend 148216 (ZET) |
| F4/80 | PE | 1:100 | Biolegend 123109 (BM8) |
| Ly-6C | APC | 1:100 | Invitrogen 17-5932-82 (HK1.4) |
| CD64 | BV605 | 1:100 | Biolegend 139323 (X54-5/7.1) |
| CD19 | Percpcy5.5 | 1:200 | Tonbo 65-0193 (ID3) |
| NK1.1 | Percpcy5.5 | 1:200 | BD 561111 (PK136) |
| Ly-6G | Percpcy5.5 | 1:200 | BD 560602 (1A8) |
| CD3 | Percpcy5.5 | 1:200 | BD 560527(17A2) |
| MHC-II | APC-cy7 | 1:500 | Biolegend 107602 (M5/114.15.2) |
| SIGLEC-H | PE-Cy7 | 1:100 | eBioscience 25-0333-82 (440c) |
| CD45 | Alexa 700 | 1:100 | BD 560566 (HI30) |

**Supplementary Table 4: CyTOF antibodies**

| Marker | Metal conjugation | Clone | Species | Manufacturer | Cat # |
|---|---|---|---|---|---|
| CD11c | 209Bi | N418 | Ms | DVS-Fluidigm | 3209005B |
| CD279 | 159Tb | 29F.1A12 | Ms | Biolegend | 135202 |
| CD45 | 147Sm | 30-F11 | Ms | DVS-Fluidigm | 3147003B |
| F4/80 | 173Yb | BM8 | Ms | Biolegend | 123102 |
| Ly-6C | 150Nd | HK1.4 | Ms | DVS-Fluidigm | 3150010B |
| Ly-6G | 141Pr | 1A8 | Ms | DVS-Fluidigm | 3141008B |
| CD80 | 171Yb | 16-10A10 | Ms, Cn | DVS-Fluidigm | 3171008B |
| CD274 | 153Eu | 10F.9G2 | Ms | Biolegend | 124303 |
| CD3e | 154Sm | 145-2C11 | Ms | Biolegend | 100302 |
| CD11b | 139La | M1/70 | Ms, Hu | Biolegend | 101249 |
| CD19 | 149Sm | 4D5 | Ms | Biolegend | 115502 |
| CD40 | 161Dy | HM40-3 | Ms, Rt | DVS-Fluidigm | 3161020B |
| CD4 | 115In | RM4-5 | Ms | Biolegend | 100506 |
| CD8a | 168Er | 53-6.7 | Ms | DVS-Fluidigm | 3168003B |

**Supplementary Table 5: Patient characteristics of PDAC-TMA dataset**

| | |
|---|---|
| **N** | 120 |
| **Mean age at diagnosis** | 64.3 |
| **Sex** | |
| M | 67 |
| F | 53 |
| **Race** | |
| White | 98 |
| Hispanic | 13 |
| Black | 5 |
| Asian | 2 |
| Other | 2 |
| **Neo-adjuvant therapy** | 0 |
| **Adjuvant chemotherapy** | |
| Y | 79 |
| N | 39 |
| Unknown | 2 |
| **Adjuvant radiation therapy** | |
| Y | 56 |
| N | 62 |
| Unknown | 2 |
| **Surgical margins** | |
| 0 | 98 |
| 1 | 22 |
| **Lymph node involvement** | |
| Y | 93 |
| N | 27 |
| **Tumor grade** | |
| Well/moderate | 85 |
| Poor | 35 |
| **Surgery** | |
| Pancreaticoduodenectomy | 104 |
| Distal pancreatectomy | 15 |
| Total pancreatectomy | 1 |
| **AJCC Stage** | |
| IA | 0 |
| IB | 1 |
| IIA | 24 |
| IIB | 91 |
| III | 1 |
| IV | 3 |
